# Telomere-driven senescence accelerates tau pathology, neuroinflammation and neurodegeneration in a tauopathy mouse model

**DOI:** 10.1101/2025.06.03.657581

**Authors:** Debora Palomares, Axelle Vanparys, Joana Jorgji, Esther Paître, Pascal Kienlen-Campard, Nuria Suelves

## Abstract

**Background:** Although the connection between aging and neurodegenerative pathologies like Alzheimer’s disease (AD) has long been recognized, the underlying pathological mechanisms remain largely unknown. Senescent brain cells build up in the brains of AD patients and a causal link has been established between senescence and AD-related tauopathy.

**Methods:** To investigate the role of cellular senescence in tau-mediated neuropathology, we crossed the Terc knockout (Terc^-/-^) senescent mouse model with the P301S tauopathy model (PS19 line). Using brain sections and protein extracts, we employed Western blot and immunostaining analyses to investigate the expression of tau-related neuropathological features within a senescent context.

**Results:** We found that the brains of 6-and 9-month-old Terc^-/-^ mice exhibit significant telomere attrition and signs of cellular senescence. Introducing a senescent phenotype in a tauopathy mouse model resulted in increased tau phosphorylation at key residues, particularly in the hippocampus. Over time, this was associated with enhanced tau truncation and aggregation. These pathological changes were accompanied by exacerbated astrocyte and microglial activation, as well as selective neuronal loss in vulnerable brain regions.

**Conclusions:** Overall, our findings place senescence as a key upstream regulator of tau pathology, suggesting that targeting senescent cells and their detrimental effects may offer promising therapeutic strategies for AD and other related tauopathies.

## Background

Aging is the main predictor for most common chronic diseases and disabilities, including many neurodegenerative diseases (NDs), of which Alzheimer’s disease (AD) is the most prevalent [1]. With the global increase in life expectancy and the consequent aging population, the prevalence of these diseases is on the rise, putting a significant strain on healthcare systems and progressively becoming a major cause of societal and economic concern worldwide. Furthermore, this burden is expected to intensify in the next few years. In 2019, the estimated global number of people living with AD was around 57 million but could reach 153 million by 2050 [2, 3]. This alarming demographic trend underscores the pressing need to unravel the precise mechanisms underlying pathological aging and its role in the onset and progression of NDs.

Substantial evidence suggests that cellular senescence, a hallmark of aging, contributes to the development of age-related pathologies [4]. Originally defined in proliferating cells, senescence refers to a state of permanent cell cycle arrest, further characterized by a complex phenotype that includes the secretion of inflammatory factors that cause detrimental effects to surrounding tissues. Telomere attrition, DNA damage, oxidative stress and oncogenic signaling are all well-known triggers of senescence, activating similar molecular pathways that induce and sustain the senescence phenotype. With age, senescent cells accumulate in tissues and promote a state of chronic inflammation. While this has been associated with tissue dysfunction and disease onset in peripheral tissues [5], its specific contribution to the development of disease in the central nervous system remains largely unexplored. A key unresolved question is whether senescence acts as an upstream trigger of neurodegeneration or appears as a consequence of the formation of brain lesions in NDs [24]. Since the precise molecular mechanisms driving senescence pathological effects are still unexplored, this remains a complex and highly debated issue which requires further investigation.

AD represents the main cause of dementia and is characterized by progressive synaptic dysfunction and neuronal loss caused by two distinct histopathological lesions: amyloid-β (Aβ)-containing plaques and neurofibrillary tangles (NFTs) of hyperphosphorylated tau. Tau pathology is also found in other neurodegenerative diseases beyond AD, collectively known as tauopathies, including corticobasal degeneration (CBD), progressive supranuclear palsy (PSP) and frontotemporal lobe degeneration (FTLD), among others [6]. While amyloid pathology is an early step in AD progression [7], tau hyperphosphorylation and aggregation occur later in the disease and closely correlate with the evolution of cognitive decline and neurodegeneration [8, 9]. Pharmacological treatments for AD are mostly symptomatic, except for very few immunotherapies against amyloid-β, which have shown only modest therapeutic benefits [10].

While a small percentage of AD cases have a known cause, linked to inherited mutations in genes involved in Aβ production, most cases are sporadic, with an unknown underlying molecular basis. However, aging is recognized as a major risk factor for both sporadic and inherited forms of AD, with an increasing number of studies suggesting that brain cellular senescence may play a role in the development of AD. Senescent neurons and glial cells have been identified in the brains of AD patients [11–17] as well as in AD-related mouse models, including tauopathy models [13, 18, 19]. Furthermore, the targeted removal of senescent cells through genetic or pharmacological approaches in a tauopathy model has been shown to alleviate tau pathology, reduce neurodegeneration and improve cognitive function [18]. Additionally, increasing evidence suggests a direct correlation between shorter telomere length and increased AD risk [20–23].

In this project, we crossed Terc knockout mice, which exhibit senescence due to telomere dysfunction, with PS19 tauopathy mice. Using this novel combined model, we investigated the involvement of senescence in the progression of tau pathology and associated pathological processes. Our results confirm the causal role of cellular senescence in tauopathy and highlight its significance in neuroinflammation and neurodegeneration. In that context, further exploring the molecular mechanisms underlying the relationship between cellular senescence and tauopathy progression could offer valuable insights for developing novel AD treatment strategies.

## Methods

### Mouse models

Heterozygous Terc knockout mice (referred to as Terc^+/-^, Strain #004132, The Jackson Laboratory), carrying a germline deletion of the telomerase RNA component Terc [25], and hemizygous Tau P301S (PS19 line) transgenic mice (referred to as Tau^Tg^, Strain #008169, The Jackson Laboratory), overexpressing the 1N4R human tau isoform with the P301S mutation found in frontotemporal dementia with parkinsonism-17 (FTDP-17) [26], were purchased and maintained on a C57BL/6 genetic background. To procure double mutant Terc^-/-^ Tau^Tg^ mice of the second (G2) and third (G3) generations, the following breeding strategy was followed. Heterozygous Terc^+/-^ mice were first crossed with hemizygous Tau^Tg^ mice to obtain Terc^+/-^ Tau^Tg^mice. These mice were then crossed with Terc^+/-^ mice to generate G1Terc^-/-^ Tau^Tg^ and G1Terc^-/-^ littermates lacking telomerase activity. G1Terc^-/-^ Tau^Tg^ and G1Terc^-/-^ mice and their offspring were subsequently crossbred to produce the later generations. Age-and sex-matched non-transgenic (wild-type, WT) and hemizygous Tau^Tg^ control littermates came from a separate breeding setup. Genotype of the mice was confirmed by polymerase chain reaction (PCR) analysis on ear tag DNA samples. All animals were housed in standard animal care facilities on a 12 h light/12 h dark cycle, with *ad libitum* access to food and water. Animals were monitored daily and sacrificed if they showed signs of pain or distress. All animal procedures were carried out in compliance with institutional and European guidelines and were approved by the Ethical Committee for Animal Welfare at UCLouvain (References: 2018/UCL/MD/38 and 2021/UCL/MD/18).

### Brain tissue sampling

Mice were anesthetized with a combination of ketamine (100 mg/kg) and medetomidine (1 mg/kg) diluted in sterile phosphate-buffered saline (PBS), followed by transcardial perfusion with ice-cold sterile PBS. The brains were removed and divided into two hemispheres. The right hemisphere was post-fixed by immersion in 4% paraformaldehyde in PBS for 24 h, cryoprotected in PBS containing 30% sucrose and 0.02% sodium azide, and subsequently frozen. The left hemisphere was further dissected into the hippocampus and cortex, which were snap frozen in dry ice and stored at –80 °C.

### Telomere length analysis by quantitative PCR

Hippocampal brain tissue obtained from mice was digested with proteinase K (100 μg/ml) in lysis buffer (100 mM NaCl, 100 mM Tris-HCl [pH 8.0], 1 mM EDTA, [pH 8.0] and 1% SDS) at 50 °C overnight. DNA was purified using the phenol/chloroform method. To measure the length of mouse telomeres, a quantitative PCR method was used as previously described [27]. Briefly, two separate quantitative PCR reactions were performed, one with telomere primers and the other with single copy gene primers. The relative telomere length (presented as T/S ratio) was calculated by comparing telomere amplification (T) to that of the single copy gene (S) using the 2^-ΔΔCT^ method [28].

### Quantitative RT-PCR

Total RNA was extracted using TriPure™ Isolation reagent (Roche) according to the manufacturer’s protocol. RNA was resuspended in DEPC-treated water and its concentration was measured (OD 260 nm) on BioSpec-nano spectrophotometer (Shimadzu Biotech). Reverse transcription (RT) was carried out with the iScript cDNA synthesis kit (Bio-Rad Laboratories) using 1 μg of total RNA in a 20 μl reaction volume. Quantitative PCR was performed for the amplification of cDNAs using the appropriate primers (Table S1) and the GoTaq® qPCR Master Mix (Promega), following manufacturer’s instructions. For negative controls, the iScript reverse transcriptase was omitted in the cDNA synthesis step. Relative quantification was calculated by the 2^-ΔΔCT^ method using *Gapdh* and *Tbp* as housekeeping genes. Results were then normalized (fold change) to the control condition.

### Protein isolation

Mouse cortical and hippocampal tissues were homogenized as follows. Briefly, tissues were sonicated in ice-cold lysis buffer containing 20 mM Tris-base (pH 8.0), 150 mM NaCl, 1% NP-40, 10% glycerol and supplemented with protease and phosphatase inhibitor cocktails (Roche). The total homogenate was centrifuged at 16,000 ×g for 20 min and the supernatant was collected as the soluble fraction and stored at –80 °C. For the analysis of tau in the insoluble fraction, the pellet was washed in ice-cold PBS and resuspended by sonication in a buffer containing 8 M urea, 5 mM EDTA, 1 mM EGTA and supplemented with protease and phosphatase inhibitor cocktails (Roche). The samples were centrifuged at 16,000 ×g for 15 min and the supernatant was collected as the insoluble fraction and stored at –80 °C.

### Western blot

Total protein concentrations in the soluble and insoluble fractions were determined using the Pierce™ BCA Protein assay (Invitrogen, Thermo Fisher Scientific). NuPAGE™ LDS Sample Buffer (Invitrogen, Thermo Fisher Scientific) and 50 mM dithiothreitol (DTT) were added. Only the soluble protein fraction samples were heated for 10 min at 70°C. 10μg to 20μg of total protein were resolved by SDS-PAGE on precast NuPAGE™ 4–12% Bis–Tris gels (Thermo Fisher Scientific) with MES-SDS running buffer (Thermo Fisher Scientific) and SeeBlue™ Plus2 pre-stained (Thermo Fisher Scientific) as standard. Samples were then transferred to 0.45μm nitrocellulose membranes (Thermo Fisher Scientific). The nitrocellulose membranes were blocked in 5% (*w/v*) non-fat powdered milk in PBS/0.1% Tween®20 (PBS-T) for 30 min-1 h at room temperature, before incubation overnight at 4 °C with primary antibodies (Table S2) diluted in PBS-T. Species-specific secondary antibodies conjugated with horseradish peroxidase (HRP, Sigma-Aldrich) diluted in PBS-T were used to bind primary antibodies prior to ECL detection (PerkinElmer). Protein band intensity was quantified using the image processing software ImageJ [29] and normalized to the protein band intensity of actin or total tau (D1M9X, Cell Signaling).

### Immunofluorescence and imaging of brain sections

Frozen right brain hemispheres were sliced into 30 μm thick coronal sections using a CryoStar™ NX50 Cryostat (Thermo Fisher Scientific) and stored at – 20 °C in cryoprotectant solution (sodium phosphate buffer containing 30% ethylene glycol and 20% glycerol) until use.

For each hemibrain, four to five free-floating sections spaced 240 μm apart containing hippocampus and entorhinal/piriform cortex – between bregma-1.94 and-2.92 based on the mouse brain atlas [30] – were selected, washed three times in 1X PBS and blocked/permeabilized for 30 min at room temperature in blocking solution containing 5% BSA, 3% normal goat serum and 0.5% Triton X-100 in 1X PBS. Subsequently, brain sections were incubated overnight at 4 °C with primary antibodies (Table S2) diluted in blocking solution. The next day, after three washes with 1X PBS, sections were incubated in the dark for 1 h at room temperature with DAPI (1:10,000) and the appropriate species-specific fluorophore-conjugated Alexa Fluor™ secondary antibodies diluted in blocking solution. Finally, sections were washed three times in 1X PBS and mounted on Superfrost™ glass slides, air dried and coverslipped using Mowiol mounting medium. Sections were stored at 4 °C. Images were captured using a ZEISS Axioscan 7 Microscope Slide Scanner with a 20X objective lens and analyzed by either ZEN (Carl Zeiss) or ImageJ [29] softwares. Briefly, we quantified the intensity and extent of the pathological phosphorylation of tau and any associated neurodegeneration and neuroinflammation by delineating the areas of interest (i.e., hippocampus, entorhinal/piriform cortex) [30] and measuring the fluorescence intensity and the percentage covered by positive signal within our regions of interest. Additionally, to study the impact of tau pathology and senescence on specific hippocampal subregions, we performed quantification of the number of neuronal cells, positive for NeuN signal. Finally, to examine the morphology of microglia, positive for Iba1 antibody, we captured images using a ZEISS LSM800 Confocal Laser Scanning Microscope with a 40X objective lens.

As negative controls, samples were processed as described in the absence of primary antibody and no signal was detected.

### Nissl staining and volume quantifications

To quantify hippocampal volume, ventricular volume and cortical thickness, four to five hemibrain sections spaced 240 μm apart largely spanning the hippocampus and entorhinal/piriform cortex, between bregma-1.94 and-2.92 based on the mouse brain atlas [30] were mounted on Superfrost™ glass slides, air dried overnight at room temperature and stained with Cresyl violet (Nissl staining). After being stained for 45 min in cresyl violet solution (0.04%), sections were dehydrated in ethanol, cleared in xylene and mounted with DPX mounting medium (Sigma-Aldrich).

For quantification of hippocampal/ventricular volumes, the brain structures were manually delineated and the areas measured. Volumes between consecutive sections were then calculated using the following formula 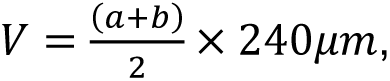 where a and b are the areas of the hippocampal/ventricular structures in each section, and 240 μm is the distance between sections. The volumes of all section pairs were added up to calculate the total hippocampal/ventricular volume [31].

### Statistical analysis

Statistical analyses were carried out using the GraphPad Prism 10 software. Normality was assessed with the Shapiro-Wilk test. A parametric test was performed if the data followed normal distribution, while a non-parametric test was used otherwise. Comparisons between two groups were made using either a parametric Student’s-t-test or a non-parametric Mann-Whitney test. When more than two groups were compared, parametric one-way ANOVA with indicated post-hoc tests or non-parametric Kruskal-Wallis were used. For correlation analyses, the non-parametric Spearman’s test was used. Significance is indicated as: ns = non-significant, \**P* < 0.05, \*\**P* < 0.01, \*\*\**P* < 0.001. The number of independent biological samples is indicated in figure legends with “n =”.

## Results

### The brains of G2Terc^-/-^ and G3Terc^-/-^ mice present telomere shortening and elevated expression of senescence markers

Given the well-established role of telomere attrition as a driver of cellular senescence, Terc knockout (Terc^-/-^) mice, which lack the RNA component of the telomerase, were used as a model of accelerated senescence. Previous data from our group demonstrated progressive telomere shortening in cortical extracts from increasing generations of Terc^-/-^ mice at 5 months of age [27]. In this study, we expanded upon these findings by evaluating telomere length in hippocampal tissue from these mice. Our results confirmed a progressive reduction in telomere length across successive generations of Terc^-/-^ mice, with significant attrition observed in the second and third generations, compared to wild-type control mice (Figure 1A).

**Figure 1.**
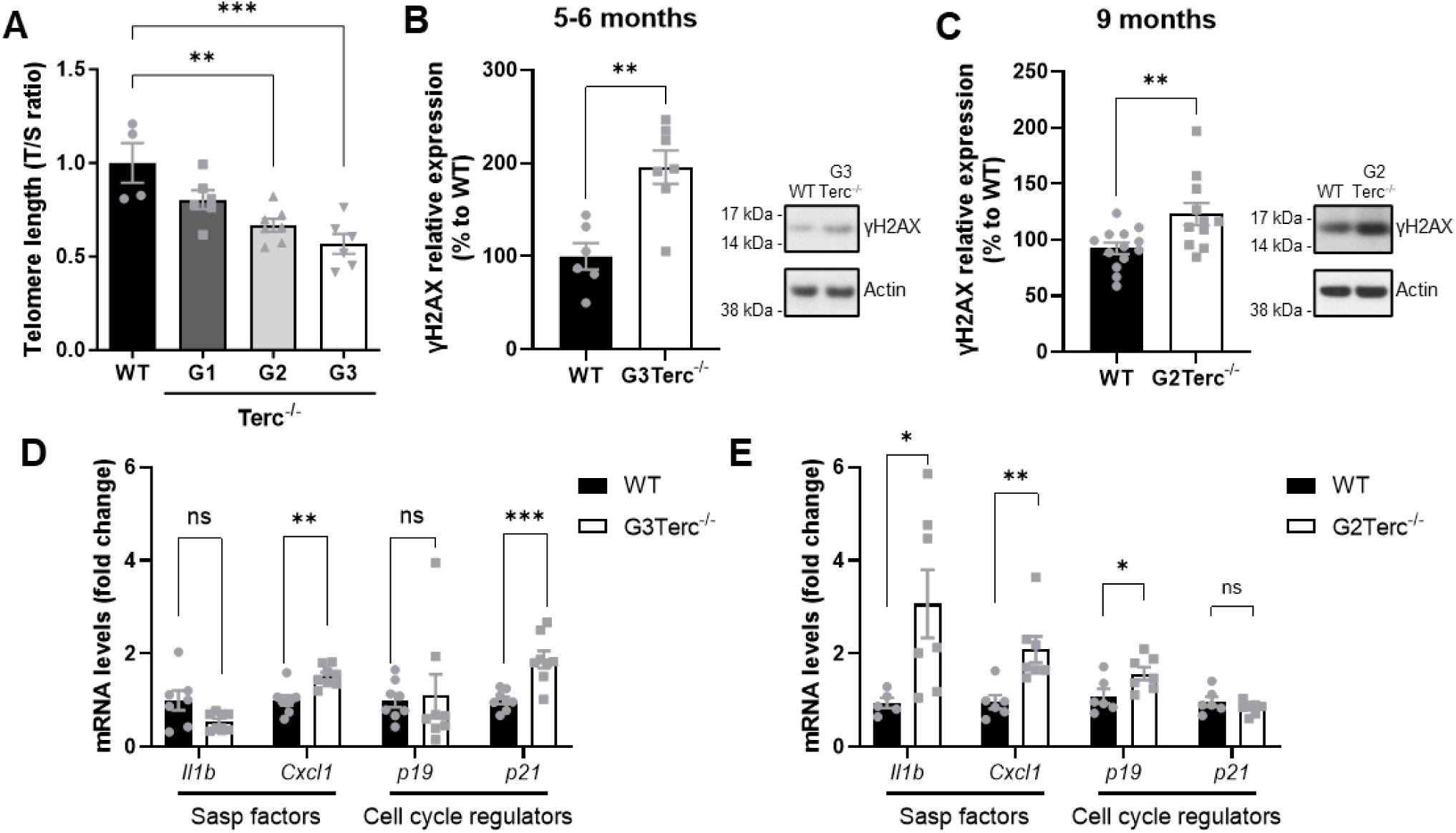
Telomere shortening in the brains of G3Terc^-/-^ and G2Terc^-/-^ mice triggers the activation of classical senescence markers. **A)** Relative telomere length detected by qPCR analysis using hippocampal DNA extracts from WT and successive generations (G1-G3) of Terc^-/-^ mice at 5 months of age. The average telomere length was calculated as the ratio (T/S) of telomere repeat copy number (T) to a single-copy gene (S = 36B4). *\*\*P* < 0.01, *\*\*\*P* < 0.001 (One-way ANOVA with Tukey’s post-hoc analysis, n = 4-7 mice/group). **B-C)** Relative levels of the DNA damage biomarker γH2AX detected by Western Blot analysis using hippocampal protein extracts from 5-6-month-old WT and G3Terc^-/-^ mice (**B**) and 9-month-old WT and G2Terc^-/-^ mice (**C**). **P < 0.01 (two-tailed Student’s *t*-test, n=6-13). **D-E)** mRNA levels of the SASP factors Il1b and Cxcl1, and the cyclin-dependent kinase inhibitors p19^Arf^ (p19) and p21^Waf1/cip1^ (p21), were measured by RT-qPCR in hippocampal extracts from 5-month-old WT and G3Terc^-/-^ mice (**D**) and 9-month-old WT and G2Terc^-/-^ mice (**E**). *\*P* < 0.05, *\*\*P* < 0.01, *\*\*\*P* < 0.001 (two-tailed Student’s *t-*test or Mann-Whitney’s test, n= 5-8 mice/group).

For our study, we selected two distinct time points from our senescent model. G3Terc^-/-^ mice at 5-6 months of age were chosen as they display a pronounced senescent phenotype, as previously confirmed in our study using cortical extracts and primary neurons derived from this generation of mice [27]. Of note, we observed a dramatic decrease in the life expectancy of G3Terc^-/-^ mice after 6-7 months of age, emphasizing the devastating effects of systemic telomere attrition. In contrast, G2Terc^-/-^ mice live longer but still show significant telomere attrition in the brain (Figure 1A). Therefore, 9-month-old G2Terc^-/-^ mice were selected to reflect an aged phenotype in a senescent context. As both models will be later used in our study to evaluate the role of senescence in tau pathology, we first validated the presence of senescence biomarkers in their brains. The evaluation of γH2AX, a phosphorylated form of histone H2AX at Ser139 that signals for double-stranded DNA breaks and is recognized as a senescence marker [32], revealed elevated levels in both G3Terc^-/-^ and G2Terc^-/-^ models (Figure 1B,C). In addition, we performed RT-qPCR analyses of *bona-fide* senescence markers, including the SASP factors Il1b and Cxcl1, and the cyclin-dependent kinase inhibitors p19^Arf^ (p19) and p21^waf1/cip1^ (p21). Increased mRNA expression of Cxcl1 and p21 was observed in 5-6-month-old G3Terc^-/-^ mice, while Il1b, Cxcl1 and p19 were upregulated in 9-month-old G2Terc^-/-^ mice (Figure 1D,E). Interestingly, the fold change in SASP factors was greater in 9-month-old G2Terc^-/-^ mice compared to 5-6-month-old G3Terc^-/-^ mice, suggesting that the aging phenotype further amplifies the secretion of pro-inflammatory molecules in a senescent context. These findings support our rationale for using both models: while G3Terc^-/-^ at 5-6 months present strong senescent features, G2Terc^-/-^ at 9 months display a combination of moderate senescence and an aged phenotype with enhanced chronic inflammation.

### Cellular senescence increases tau phosphorylation, cleavage and aggregation in PS19 mice

To explore the role of cellular senescence in the onset and development of tau pathology, we generated a novel combined mouse model by crossing Terc knockout mice with the PS19 tauopathy model, hereafter referred to as Tau^Tg^, a transgenic mouse that overexpresses the 1N4R human tau isoform with the P301S mutation found in familial FTLD. This widely used and well-characterized model exhibits widespread aberrant tau phosphorylation and aggregation by 6 months of age and significant neuronal loss by 9-12 months of age, with a particularly strong effect on the hippocampus and the entorhinal cortex [26, 33]. To assess the impact of senescence at two different stages of pathology in Tau^Tg^ mice, we selected adult mice at 6 months and relatively aged mice at 9 months of age. For each time point, four different groups of mice were obtained. At 6 months of age, we collected samples from WT, G3Terc^-/-^, Tau^Tg^, and G3Terc^-/-^ Tau^Tg^ mice. As mentioned above, G3Terc^-/-^ mice in our facilities hardly survived past 6-7 months, so we also collected samples from WT, G2Terc^-/-^, Tau^Tg^, and G2Terc^-/-^ Tau^Tg^ mice at 9 months of age (Figure S1).

To assess pathological tau phosphorylation, we first performed immunofluorescence (IF) staining analyses on coronal brain sections from all groups of mice (Figure 2). The AT8 antibody, which detects tau phosphorylated at Ser202 and Thr205, is a well-established marker of tau pathology [34, 35]. Co-labeling with an anti-NeuN antibody enabled the visualization of pathological tau in neuronal somas. AT8 signal was negligible in 6-month-old WT and G3Terc^-/-^ mice, as well as in 9-month-old WT and G2Terc^-/-^ mice (Figure 2A,B), whereas strong AT8 labeling was detected in hippocampal and cortical regions from tauopathy and combined senescence-tauopathy models. Image quantification revealed an increase in AT8 signal intensity and signal-covered area in 6-month-old G3Terc^-/-^ Tau^Tg^ compared to Tau^Tg^mice, which was even more exacerbated in 9-month-old G2Terc^-/-^ Tau^Tg^ mice. Further characterization in 9-month-old G2Terc^-/-^ Tau^Tg^ mice revealed a regional increase in AT8 staining within the hippocampus and the piriform/entorhinal cortex, suggesting increased vulnerability of these regions to the senescent context (Figure 2C).

**Figure 2.**
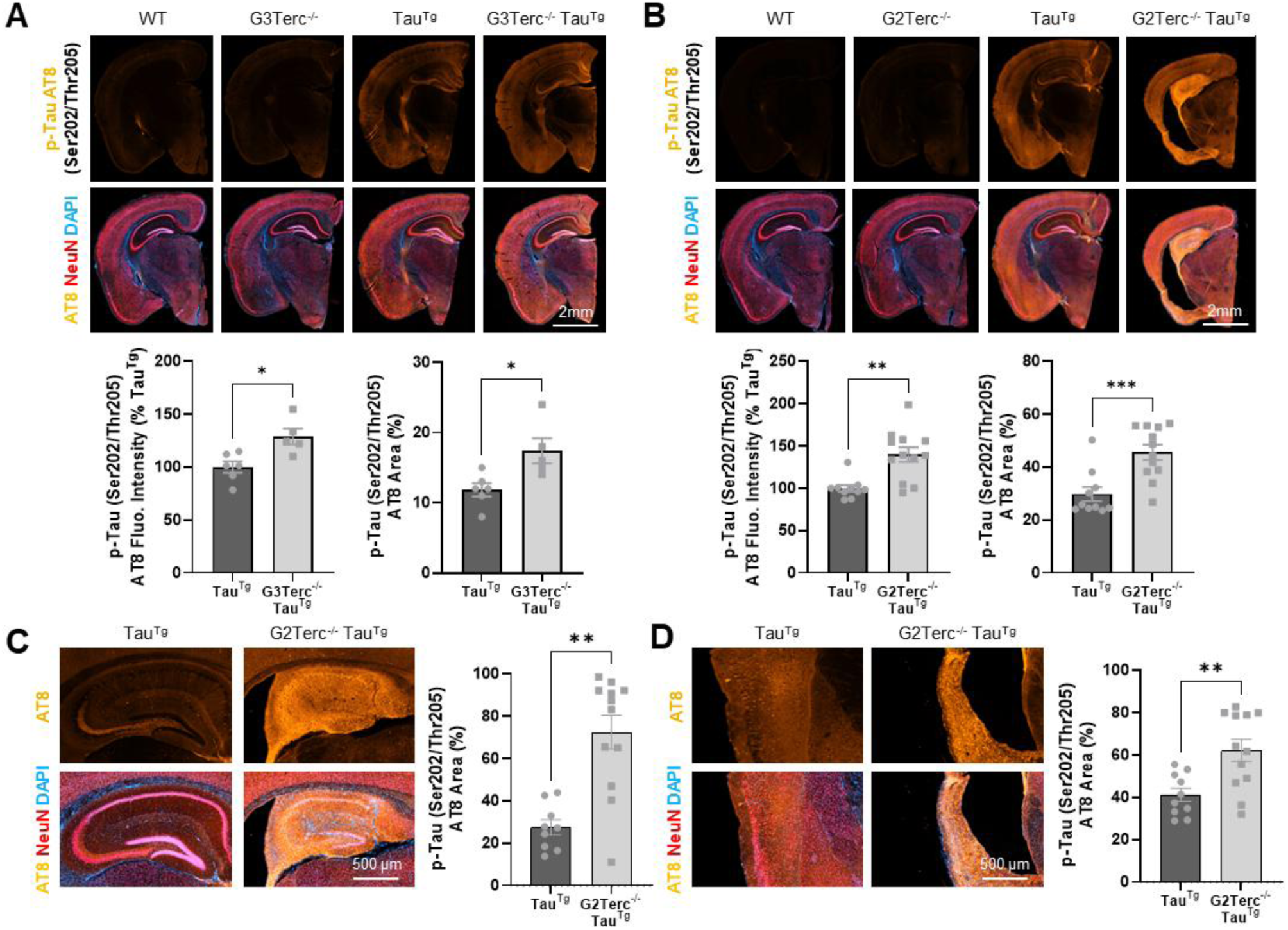
Increase of pathologically phosphorylated tau in a senescent context. Immunofluorescence analysis of pathologically phosphorylated tau (Ser202/Thr205 – AT8, orange), co-localized with the neuronal marker NeuN (red) and DAPI (blue), in brain sections from 6-month-old WT, G3Terc^-/-^, Tau^Tg^ and G3Terc^-/-^ Tau^Tg^ mice (**A**) and 9-month-old WT, G2Terc^-/-^, Tau^Tg^ and G2Terc^-/-^ Tau^Tg^ mice **(B-D)**. Magnification insets of the hippocampal region **(C)** and piriform/entorhinal cortex **(D)** are shown for immunostainings displayed in B. The relative fluorescence intensity and the percentage of area covered by AT8 is calculated for the whole brain section (A, B) and the percentage of area covered by AT8 is calculated for the hippocampal region (C) and entorhinal/piriform cortex (D). \**P* < 0.05, \*\**P* < 0.01, \*\**P* < 0.001 (Student’s *t-*test, n=5-12 mice/group).

To conduct a more in-depth evaluation of the effect of senescence on tau phosphorylation status, we performed WB analyses of several tau epitopes known to be hyperphosphorylated in pathological conditions: Thr181, Ser202/Thr205 (AT8), Thr231, Ser262, Ser396/Ser404 (PHF-1) and Ser422 [36]. In the adult Tau^Tg^ mouse brain, endogenous murine tau, which exists primarily as the 0N4R isoform [37, 38], coexists with the human 1N4R tau isoform. This isoform difference results in two bands of different molecular weights in WB analyses using anti-tau antibodies, enabling the discrimination between mouse and human tau and the assessment of their phosphorylation status. However, it has been reported that commonly used phospho-tau antibodies, when evaluated by WB in mouse brain samples, display non-specific bands at ∼50 kDa, the molecular size of murine tau. These bands can still be observed in samples from Tau knockout mice and therefore cannot be attributed to phosphorylated endogenous tau [39]. For this reason, we focused our evaluation on the phosphorylation status of human tau, using only the tauopathy and combined senescence-tauopathy models at 6 and 9 months of age. Our analyses revealed a marked increase in tau phosphorylation at Thr181 and Ser202/Thr205 (AT8) in both soluble and insoluble protein fractions, and at Thr231 and Ser396/Ser404 (PHF-1) only in the soluble fraction, in the hippocampus of 6-month-old G3Terc^-/-^ Tau^Tg^ (Figure 3A). No changes in Ser262 and Ser422 residues were found. Unlike the hippocampus, the cortical region of these mice showed no changes at any of the evaluated phosphorylation sites (Figure 3B). Evaluation of total tau levels revealed no significant differences in either soluble or insoluble protein fractions (Figure S2). At a later timepoint in the disease progression (9 months), all the evaluated tau phosphorylation residues were significantly increased in the insoluble protein fraction of the hippocampus in G2Terc^-/-^ Tau^Tg^ mice (Figure 3C), indicating that tau pathology is aggravated under a senescent context. Notably, the fold change differences were greater at 9 months compared to 6 months. Tau phosphorylation at Thr181 and Ser202/Thr205 (AT8) was still increased in the soluble fraction. The cortical region was again relatively spared, although a significant increase was found in phosphorylated tau at Thr231 and Ser422 in the soluble fraction and an almost significant trend towards an increase was found in Ser202/Thr205 (AT8) and Ser422 sites in the insoluble fraction (Figure 3D). Additionally, total tau levels were significantly elevated in soluble and insoluble protein fractions of these mice (Figure S2). Overall, our data suggests that the senescent context induces site-specific hyperphosphorylation of tau in a region-dependent manner, eventually leading to an increase in tau accumulation and deposition.

**Figure 3.**
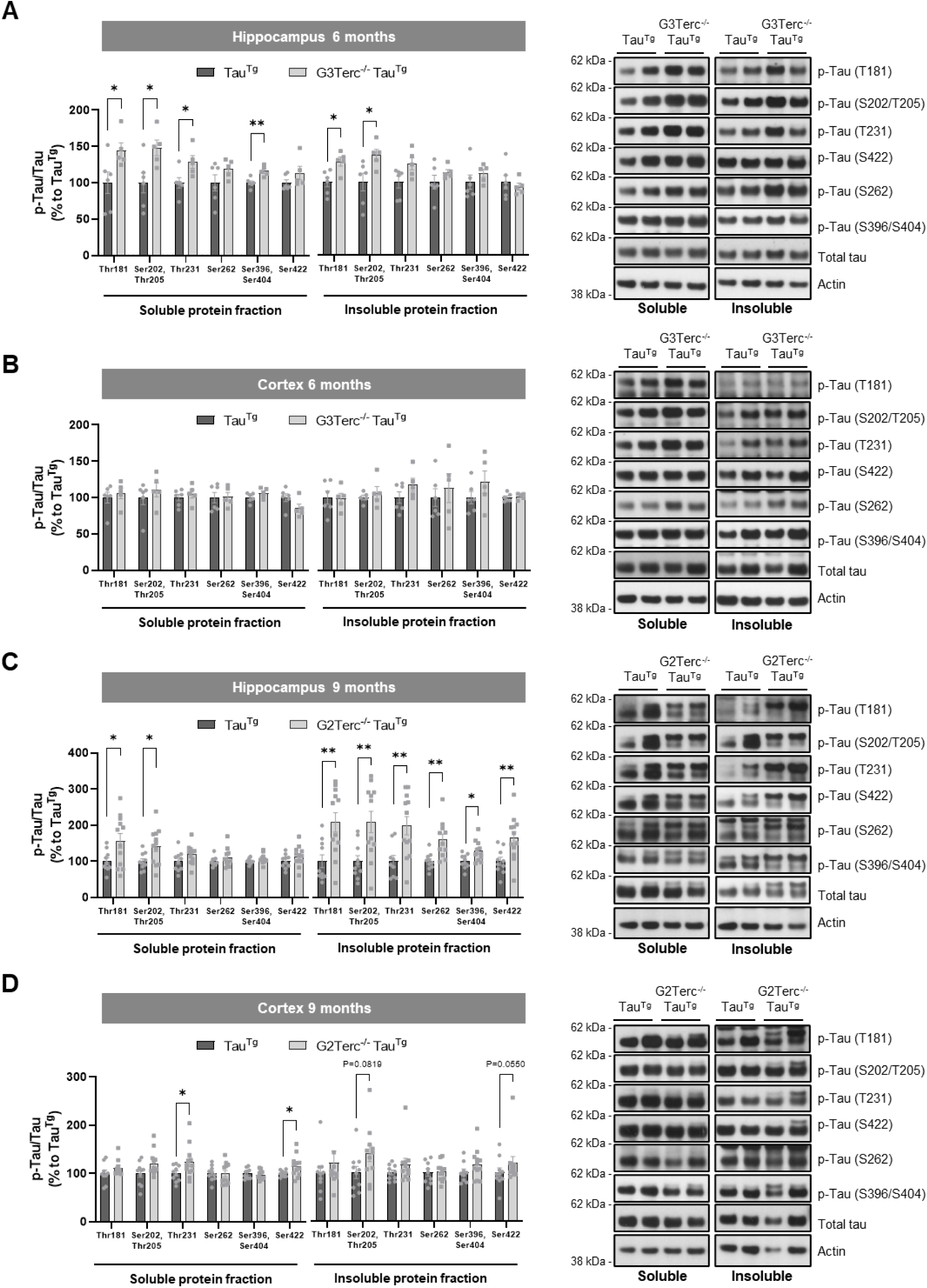
Senescence induces site-specific hyperphosphorylation of tau in PS19 mice. WB analysis of phosphorylated tau (Thr181, Ser202/Thr205 – AT8, Thr231, Ser262, Ser396/Ser404 – PHF-1 and Ser422) normalized to total tau in soluble and insoluble protein extracts from 6-month-old Tau^Tg^ and G3Terc^-/-^ Tau^Tg^ hippocampal **(A)** and cortical **(B)** samples and from 9-month-old Tau^Tg^ and G2Terc^-/-^ Tau^Tg^ hippocampal **(C)** and cortical **(D)** samples. \**P* < 0.05, *\*\*P* < 0.01 (Student’s *t-*test or Mann-Whitney’s test, n= 5-12 mice/group).

To further investigate the molecular mechanisms underlying these effects, we assessed several major tau kinases, including AMP-activated protein kinase (AMPK), glycogen synthase kinase 3β (GSK-3β), mitogen-activated protein kinases ERK1/2 and protein kinase B (also known as AKT), as well as the primary tau phosphatase, protein phosphatase 2A (PP2A) (Figure S3). Telomere attrition-induced senescence had no major impact on the protein levels and phosphorylation status of most tau kinases and phosphatases at 6 months of age, although a significant decrease in the active form of AMPK (p-AMPK) was observed in G3Terc^-/-^ Tau^Tg^ mice compared to WT controls. This suggests that the early effects of senescence on tau phosphorylation might not be driven by the over-activation of tau kinases or inactivation of tau phosphatases. In contrast, at 9 months of age (Figure S3), p-ERK becomes hyperactive in the hippocampi of Tau^Tg^ mice, with further enhancement caused by the senescence context, while p-AKT was found significantly increased in G2Terc^-/-^ Tau^Tg^ mice. Additionally, a significant decrease in p-AMPK was found in Tau^Tg^ mice, becoming further inactivated in G2Terc^-/-^ Tau^Tg^. These results suggest that advanced tau pathology disrupts the activity of AMPK, ERK1/2 and AKT kinases, and that this dysregulation might be accelerated by the senescent environment.

Since our data showed that telomere-driven senescence promotes tau phosphorylation, particularly in the hippocampus, we sought to determine whether it also modulates tau cleavage and aggregation in this region. WB analysis using a total tau antibody (D1M9X clone) on soluble and insoluble hippocampal protein fractions of 9-month-old mice revealed several bands below 50 kDa in Tau^Tg^ and G2Terc^-/-^ Tau^Tg^ mice, which were absent in WT and G2Terc^-/-^ samples (Figure S4). This suggests that the observed bands correspond to specific fragments of the tau transgene, as previously indicated [40, 41]. Tau truncation by different proteases in the brains of AD patients and tauopathy models has been involved in tau aggregation and neurotoxicity [42, 43]. Our WB evaluation of truncated tau in hippocampal samples showed a trend towards an increase in the soluble fraction and a significant increase in the insoluble fraction in G2Terc^-/-^ Tau^Tg^ mice compared to Tau^Tg^ mice (Figure 4). Similarly, the total tau antibody in soluble and insoluble protein fractions from 9 months-old Tau^Tg^ and G2terc^-/-^ Tau^Tg^ revealed several bands of high-molecular weights, indicative of tau oligomers [40, 41]. Again, these bands were absent in WT and G2Terc^-/-^ samples (Figure S4). WB analyses demonstrated increased levels of oligomeric tau species in the hippocampus of G2Terc^-/-^ Tau^Tg^ mice compared to Tau^Tg^ mice, in both soluble and insoluble protein fractions (Figure 4). These data suggest that telomere-induced senescence increases tau truncation and aggregation in relatively aged PS19 mice.

**Figure 4.**
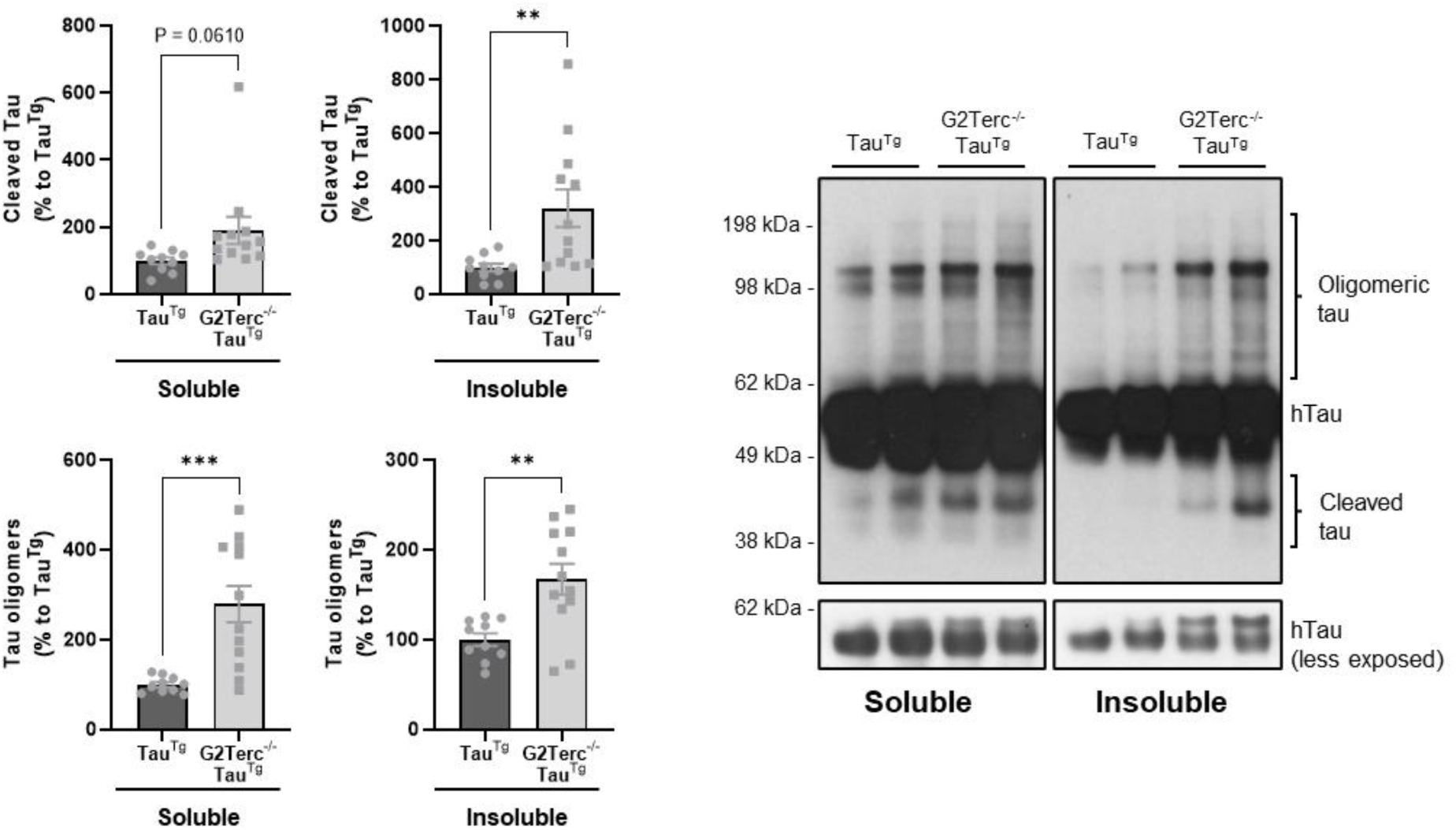
Senescence exacerbates truncated and aggregated tau. WB evaluation of oligomeric and truncated Tau levels in soluble and insoluble protein extracts from 9-month-old Tau^Tg^ and G2Terc^-/-^ Tau^Tg^ mouse hippocampi, using a total tau antibody (D1M9X clone). \*\**P* < 0.01, *\*\*\*P* < 0.001 (Student’s *t-*test or Mann-Whitney’s test, n= 10-12 mice/group).

### Cellular senescence enhances glial activation in the presence of tauopathy

It is currently well established that glial activation and inflammation are both processes heavily involved in the development and evolution of NDs, including AD [44, 45]. Moreover, it is believed astrocyte and microglial alterations associated with aging can further contribute to the process of neurodegeneration [46–48]. Given that our results revealed strong tau pathology in the combined senescence-tauopathy model at 9 months of age, we initially assessed the presence of astrogliosis and microgliosis across all mouse groups by performing WB analyses of GFAP and Iba1 protein, which are specific markers of astrocytes and microglia, respectively (Figure 5A). Although GFAP levels showed a non-significant increasing trend in Tau^Tg^ mice compared to WT mice in the hippocampal region, the addition of a senescent context strongly upregulated GFAP levels. Similarly, cortical samples of G2terc^-/-^ Tau^Tg^ mice displayed raised GFAP levels compared to WT and Tau^Tg^ mice. Iba1 levels displayed similar changes, with G2terc^-/-^ Tau^Tg^ hippocampal and cortical samples revealing increased Iba1 levels. Notably, the change in Iba1 levels was more pronounced in the hippocampus compared to the cortex, suggesting stronger microglial activation in this region. Immunofluorescence staining for Iba1 and GFAP confirmed the activation of astrocytes and microglia in the hippocampus of 9-month-old G2terc^-/-^ Tau^Tg^ mice (Figure 5B), showing a significant increase in the biomarker-positive area for both glial cell types, when compared to WT but also to Tau^Tg^ mice. An in-depth evaluation of the morphology of Iba1-positive cells revealed that microglial activation in the hippocampus of 9-month-old G2terc^-/-^ Tau^Tg^ mice was accompanied by the presence of dystrophic microglia [49], characterized by beading and apparent fragmentation of their projections (Figure S5). Interestingly, this phenotype was not observed in Tau^Tg^ mice, despite the aforementioned increase in Iba1 levels. Altogether, these findings show that telomere-induced brain senescence, typically associated with low-grade chronic inflammation, results in the abnormal activation and proliferation of astroglia and microglia over time, particularly within a pathological background.

**Figure 5.**
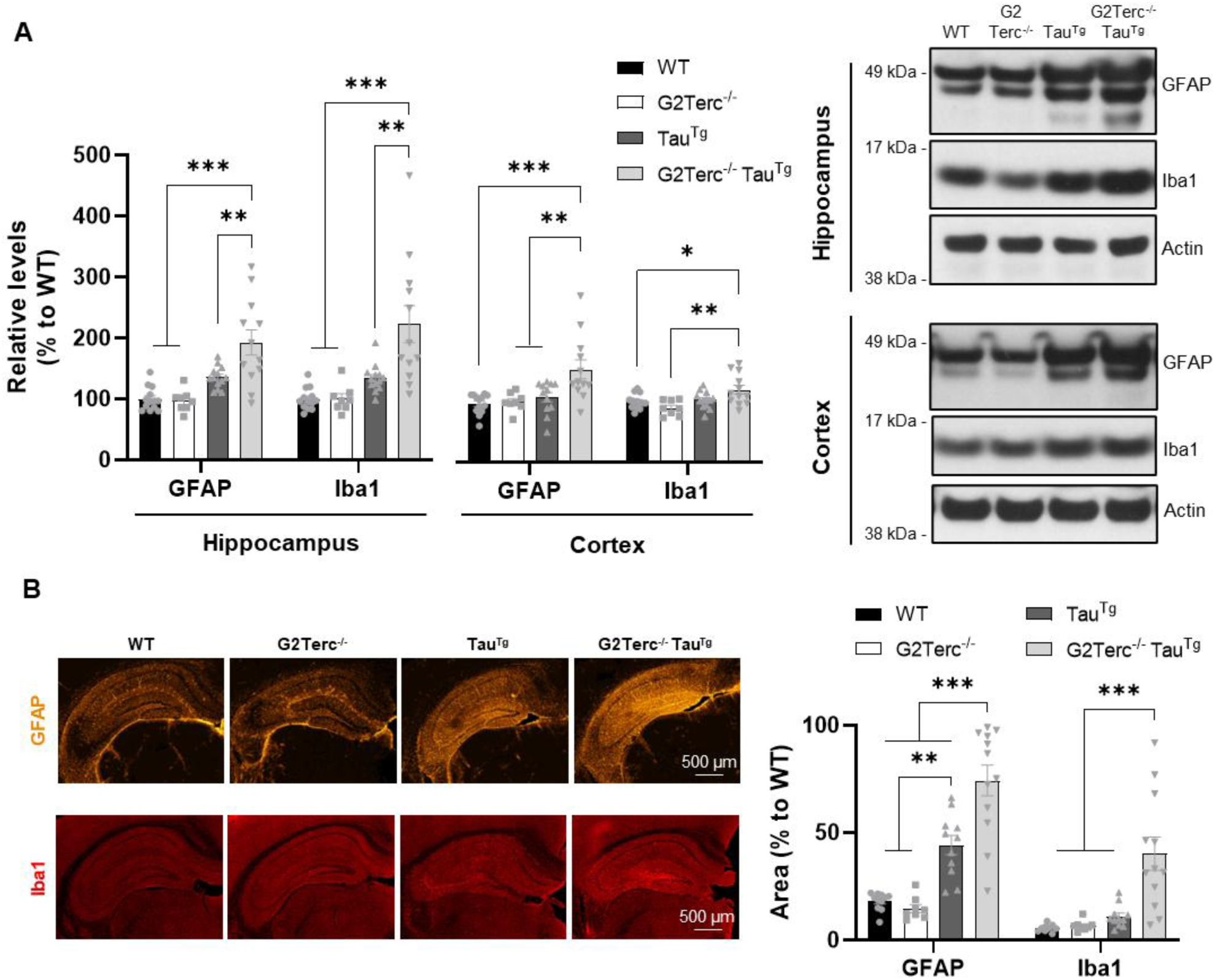
Senescence increases glial activation within a tauopathy context. **A)** WB evaluation of GFAP and Iba1 expression in hippocampal and cortical protein extracts from 9-month-old WT, G2Terc^-/-^, Tau^Tg^ and G2Terc^-/-^ Tau^Tg^ mice. \**P* < 0.05, \*\**P* < 0.01, \*\*\**P* < 0.001 (One-way ANOVA with Tukey’s post-hoc analysis, n= 8-13 mice/group). **B)** Immunofluorescence analysis of GFAP (orange) and Iba1 (red) in the hippocampal region of brain sections from 9-month-old WT, G2Terc^-/-^, Tau^Tg^ and G2Terc^-/-^ Tau^Tg^ mice. Histograms represent the percentage of staining area. \**P* < 0.05, \*\**P* < 0.01, \*\*\**P* < 0.001 (One-way ANOVA with Tukey’s post-hoc analysis, n= 8-13 mice/group).

### Senescence-induced tauopathy in PS19 mice correlates with neurodegeneration

Synaptic impairment is a pathological hallmark of neurodegeneration associated with AD and thought to precede neuronal loss in humans and in preclinical models of the disease, including the PS19 tauopathy mouse model [26, 50]. It is believed that reactive astroglia and activated microglia play an active role in the process of synaptic loss in tau pathology via enhanced synaptic phagocytosis [51, 52]. To assess the impact of tau hyperphosphorylation and neuroinflammation in neurodegeneration, we first examined the levels of well-established synaptic markers via WB. At 6 months of age, we did not observe significant changes in the protein levels of NMDAR2B, GluR1, PSD95 and synaptophysin (Figure 6A). We then examined the same synaptic markers in the hippocampus of 9-month-old G2terc^-/-^ Tau^Tg^ mice, and we observed a clear decreasing trend, which was significant for GluR1 (Figure 6B). Next, we assessed neuronal degeneration using Nissl staining and NeuN immunoreactivity analysis. While increased tau phosphorylation was observed in the hippocampus of 6-month-old G3Terc^-/-^ Tau^Tg^ mice (Figure 2A), our results demonstrated overall preservation of neuronal integrity at that age, with no detectable changes in NeuN protein levels (Figure S6A) and no gross morphological changes (Figure S6B). In contrast, G2Terc^-/-^ Tau^Tg^ brains at 9 months of age exhibited important brain atrophy, with a significant reduction in hippocampal volume, as well as a shrinkage of the piriform/entorhinal cortex (Figure 7A), compared to either WT or Tau^Tg^ controls. In contrast, cortical thickness at the level of the somatosensory area was unchanged between groups. In addition, G2Terc^-/-^ Tau^Tg^ mice also displayed marked enlargement of the lateral ventricles, which is frequently associated with hippocampal and cortical atrophies. Moreover, NeuN protein levels of 9-month-old G2Terc^-/-^ Tau^Tg^ hippocampal extracts showed a significant decrease when compared with WT, G2Terc^-/-^ and Tau^Tg^ samples, which was not present in cortical extracts (Figure 7B). Detailed examination of specific hippocampal subregions of G2Terc^-/-^ Tau^Tg^ mice revealed significant neuronal loss in the CA1, CA3 and DG (Figure 7C). Our findings further revealed that hyperphosphorylated tau (AT8) and markers of neuroinflammation (GFAP/Iba1) are negatively correlated with hippocampal volume in the brains of 9-month-old Tau^Tg^ and G2Terc^-/-^ Tau^Tg^ mice (Figure S7), suggesting a potential causal relationship. Together, our results suggest that increased tauopathy caused by senescence activation in PS19 mice leads to region-specific neuronal vulnerability.

**Figure 6.**
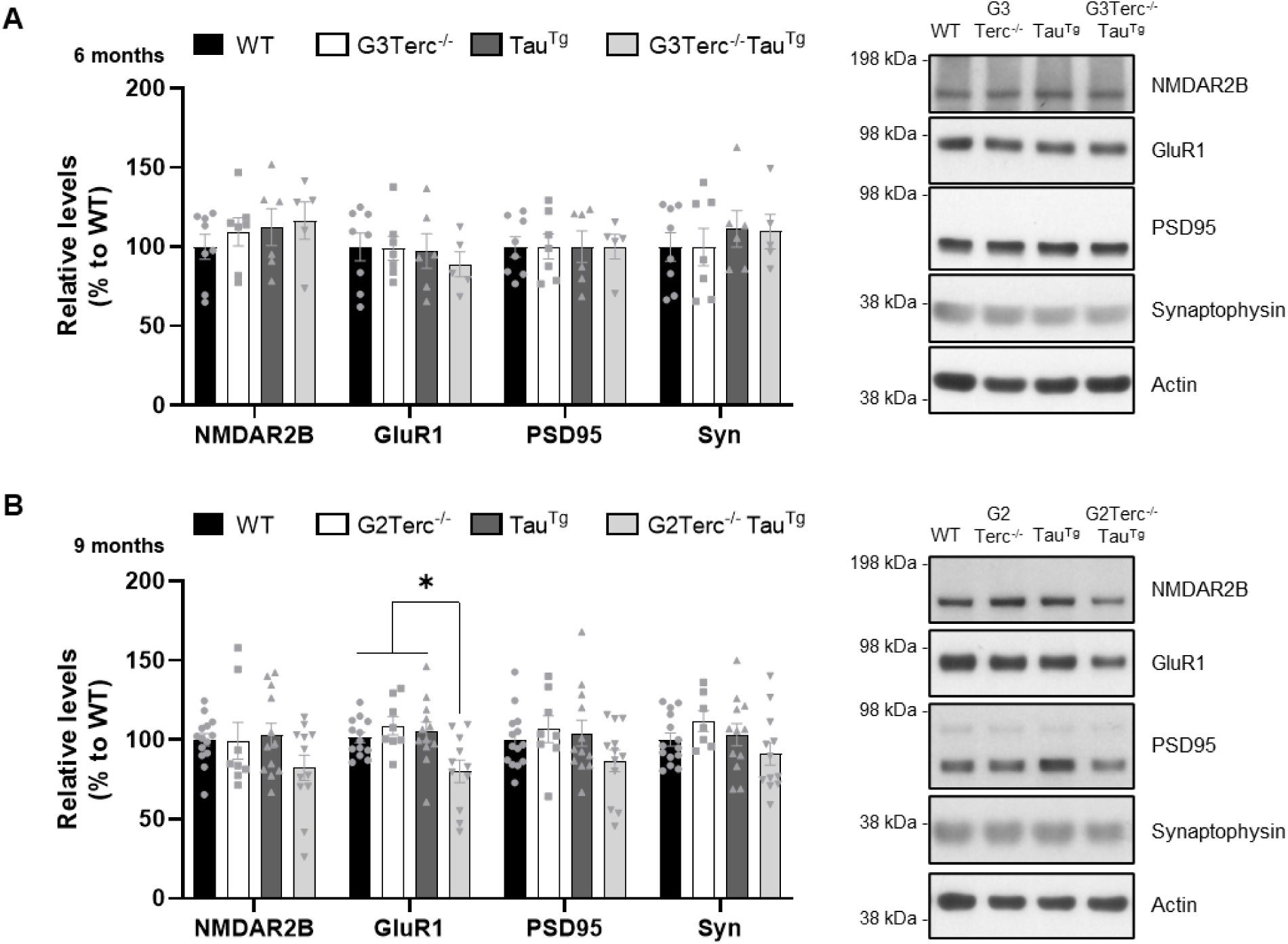
Senescence slightly impairs synaptic marker expression within a tauopathy context. WB evaluation of relative levels of the synaptic markers NMDAR2B, GluR1, PSD95 and synaptophysin in hippocampal protein extracts from 6-month-old WT, G3Terc^-/-^, Tau^Tg^ and G3Terc^-/-^ Tau^Tg^ mice (**A**) and 9-month-old WT, G2Terc^-/-^, Tau^Tg^ and G2Terc^-/-^ Tau^Tg^ mice (**B**). \**P* < 0.05 (One-way ANOVA with Tukey’s post-hoc analysis, n= 5-13 mice/group).

**Figure 7.**
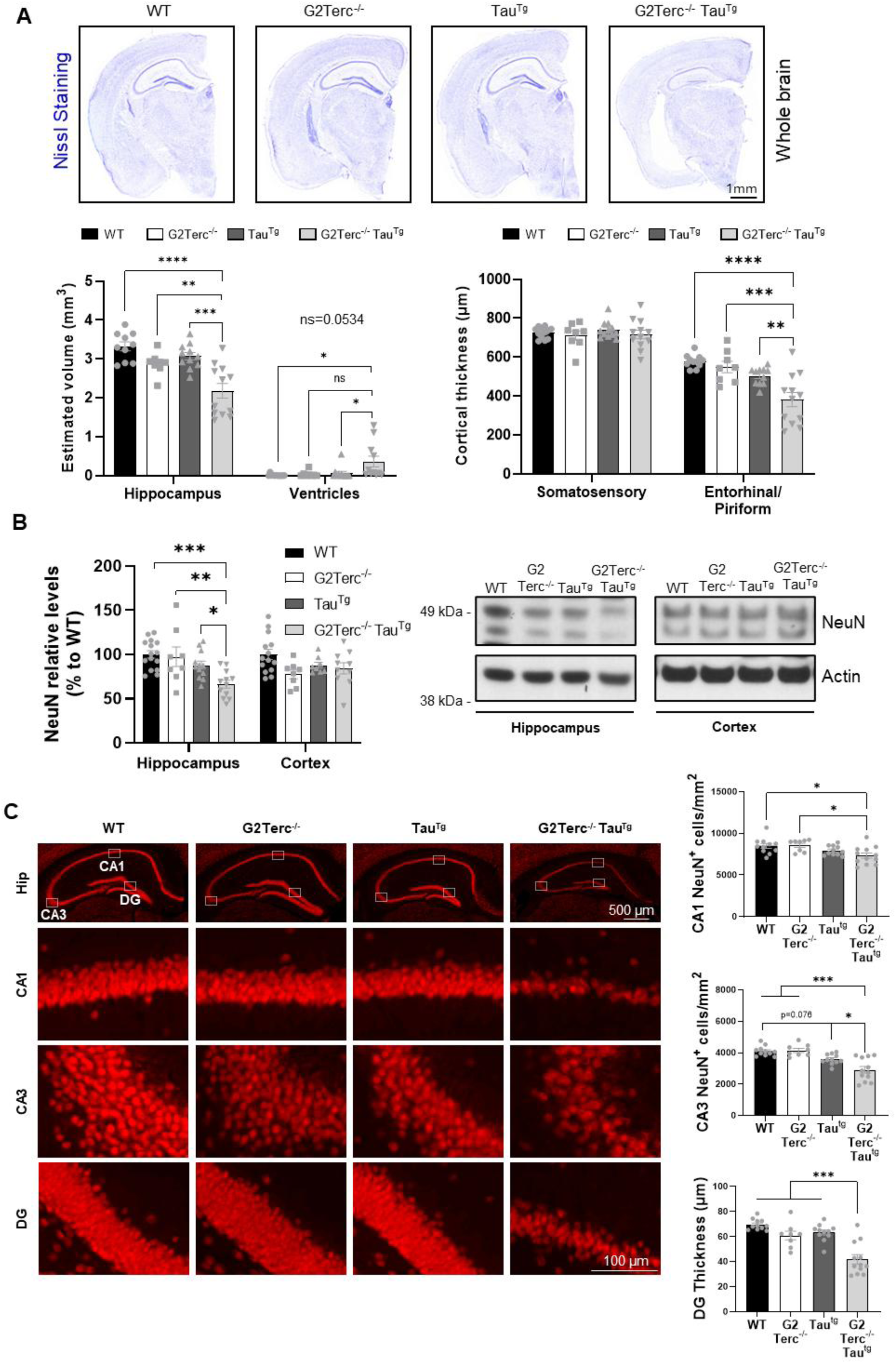
Senescence induces neurodegeneration in vulnerable brain regions of tauopathy mice. **A)** Representative Nissl staining images from 9-month-old WT, G2Terc^-/-^, Tau^Tg^ and G2Terc^-/-^ Tau^Tg^ mice. Quantitative analyses include measurements of lateral ventricle and hippocampal volumes, as well as thickness of the somatosensory cortex and entorhinal/piriform cortices. **B)** WB evaluation of NeuN expression in hippocampal and cortical extracts from 9-month-old WT, G2Terc^-/-^, Tau^Tg^ and G2Terc^-/-^ Tau^Tg^ mice. \**P* < 0.05, \*\**P* < 0.01, \*\**P* < 0.001 (One-way ANOVA with Tukey’s post-hoc analysis, n= 8-13 mice/group). **C**) NeuN immunofluorescence analysis in 9-month-old WT, G2Terc^-/-^, Tau^Tg^ and G2Terc^-/-^ Tau^Tg^ mice. Histograms show the density of NeuN-positive cells in the CA1 and CA3 areas, as well as the thickness of the DG granule cell layer. *P* < 0.05, \*\**P* < 0.001 (One-way ANOVA with Tukey’s post-hoc analysis, n= 8-13 mice/group).

## Discussion

Given that cellular senescence is closely associated with pathological aging, its role in age-related NDs like AD has attracted increasing attention in recent years [53]. Senescent neurons and glial cells have been found in AD brains [11–15], and considerable efforts have been made to evaluate their contribution to disease pathogenesis. Since telomere attrition is a primary driver of senescence, the telomerase-deficient Terc knockout mice [25] provide an excellent model for studying the role of brain senescence in the onset and progression of AD pathology.

Prior work from our group demonstrated that the senescence context present in Terc knockout mouse brains can drive toxic intraneuronal Aβ accumulation upon overexpression of human APP carrying familial AD (FAD) mutations [27]. In this project, we explored the potential contribution of senescence on another hallmark of AD pathology, tau aggregation, by crossing Terc knockout mice with the PS19 tauopathy mouse model. We found that the introduction of a senescent phenotype increased tau phosphorylation and aggregation in specific brain regions, ultimately resulting in excessive neuroinflammation and neuronal loss.

While mounting evidence suggests that AD-related features, particularly tau aggregation, can trigger cellular senescence [11, 15], the inverse relationship seems less well-established [24]. Our study reinforces the idea that senescence is an upstream regulator of the tauopathy process. This finding aligns itself with recent studies exploring the causal role of cellular senescence in tau-related pathology, where it was shown that the pharmacological or genetic elimination of senescent astrocytes and microglial cells in a tauopathy mouse model reduced tau aggregation and associated cognitive deficits [18]. Based on the existing experimental evidence [11, 15], we propose that the initial effects of senescence can become progressively amplified through a negative feedback loop as the disease advances, accelerating neurodegeneration and cognitive decline over time.

Controversial data exists regarding the presence of telomere erosion in brain tissue from AD patients, mainly due to technical limitations inherent to postmortem samples [20]. However, it seems well established that cellular senescence occurs in AD and is more prevalent than in non-affected individuals [11–17]. In fact, oxidative stress and DNA damage, which are consistently elevated in aged and AD brains, are also well-known inducers of cell senescence and are prevalent in postmitotic cells like neurons [54]. The Terc knockout mouse model of accelerated senescence used in our study is designed to create the ideal conditions for inducing senescence in a controlled, reproducible and time-efficient manner. Indeed, our findings confirm that the presence of significantly shorter telomeres in the mouse brain exacerbates classical senescent features, including the upregulation of cell cycle inhibitors and SASP factors. Telomerase inactivity is also translated into brain DNA damage, as demonstrated by increased γH2AX levels in G2Terc^-/-^ and G3Terc^-/-^ mouse hippocampi. A recent study demonstrated that the specific accumulation of undigested damaged DNA in the cytosol of neurons [55] or microglia [56] due to DNAse II inactivation can lead to increased tau phosphorylation and neurodegeneration.

Our study provides critical insights into the accumulating effect of chronic senescence on tau phosphorylation and aggregation. We found that senescence initially affects specific phosphorylation residues on tau protein, with Thr181, Ser202 and Thr205 being particularly impacted during the early phases of tauopathy progression. As the disease advances in relatively aged G2Terc^-/-^ Tau^Tg^ mice, senescence leads to a shift from detergent-soluble to detergent-insoluble phosphorylated tau forms, affecting numerous residues. This suggests that senescence plays a crucial role in the progression of tau pathology, further confirmed by the observed increase in high-molecular-weight tau species, indicative of tau oligomers, both in soluble and insoluble protein fractions. These tau oligomers are in turn associated with enhanced neurotoxicity, prompting worsened synaptic dysfunction and neurodegeneration [57, 58]. While our results showed that kinases such as AMPK, GSK3β, ERK1/2 or AKT, along with protein phosphatase 2, may not be involved in the early effects of senescence on tau phosphorylation, further investigation is needed to determine the underlying molecular mechanisms. In contrast, during advanced disease stages, our findings highlight that senescence activates the kinases ERK1/2 and AKT, which regulate tau phosphorylation [59, 60]. In addition, several cellular pathways known to be altered during senescence could have contributed to the effects we observed. Results from our previous study suggest that the autophagy flux is altered in the brains of Terc knockout mice [27], and a direct link between autophagy defects and increased tau phosphorylation and aggregation has been made [61]. While hyperphosphorylated tau is degraded by either the proteasome or the autophagy-lysosome systems in physiological conditions [62, 63], under pathological circumstances such as senescence, the increase in hyperphosphorylated tau cannot be efficiently cleared by these pathways [64]. Similarly, Terc knockout mice also present mitochondrial dysfunction and increased oxidative stress [65], two interrelated features that have been correlated with increased tau phosphorylation [66–69].

Aside from tau phosphorylation, the proteolytic processing of tau has also been involved in promoting its aggregation [42, 43]. Tau can be cleaved by multiple proteases, including calpains, caspases and endopeptidases, and the resulting fragments can be found in AD and other tauopathies at early disease stages and correlate with the degree of cognitive dysfunction [70–73]. Additionally, truncated tau has been shown to play a key role in promoting tau aggregation and inducing neurodegeneration, as demonstrated in animal models [74–76] and in vitro systems [43, 77, 78]. Our data demonstrates that telomere-induced senescence accelerates tau cleavage in PS19 mice, potentially contributing to the increased tau aggregation and neurodegeneration observed in our study. Although the precise molecular mechanisms behind this effect were not investigated, it is likely that the reported increase in calpain/caspase-3 activities during DNA damage contributes to tau fragmentation [79, 80].

While the hippocampal region appeared particularly vulnerable to the impact of senescence on tauopathy induction, the cortical region of these mice remained relatively unaffected, particularly at early stages of the disease. WB analyses revealed no significant changes in cortical tau phosphorylation at 6 months of age, while 9-month-old senescent mice displayed increased tau phosphorylation at only a few residues in the soluble protein fraction. Similarly, volume reduction and neuronal loss were found in G2Terc^-/-^ Tau^Tg^ hippocampi, but cortical extracts showed unaltered NeuN levels, and we found no thinning of the somatosensory cortical area. However, not all cortical subregions seemed equally spared. Immunostaining analyses revealed a specific vulnerability of the piriform/entorhinal cortical regions, displaying exacerbated tau phosphorylation and atrophy. This distinct vulnerability to tau pathology in the brain across different regions is also present in AD, where brain structure imaging and post-mortem examinations suggest that some areas remain relatively spared until the late stages of the pathology [81, 82].

Neuroinflammation has been shown to play a dual role in neurodegenerative conditions: while it facilitates repair during acute stages, it can promote neuronal loss and aggravate disease progression in chronic stages [45]. Our findings demonstrate that the senescence-impact on tau pathology induces neuroinflammation by increasing the activation of astrocytes and microglia in the hippocampi of old G2Terc^-/-^ Tau^Tg^ mice. On the other hand, despite a significant transcriptional upregulation of inflammatory markers in the hippocampi of our senescent model (i.e., *Il1b, Cxcl1*), levels of specific astrocyte and microglia markers, as well as the total area they occupied in this region, were not altered in the absence of tauopathy. This suggests a lack of glial activation in non-pathological contexts, consistent with our previous data [27]. Our results also indicate that microglial/astrocytic activation might be a direct consequence of tau pathology, as previously suggested [83], rather than merely a result of senescence activation. Indeed, previous studies have shown that microglia and astrocytes from Terc knockout brains do not display major functional alterations, unless in the presence of pathological protein aggregates or inflammatory stimuli [84–86]. Notably, dystrophic microglia were observed in the hippocampus of 9-month-old G2terc^-/-^ Tau^Tg^ mice, but not in Tau^Tg^ mice, suggesting that the combination of senescence and tauopathy has a distinct and more detrimental impact on microglia than tauopathy alone.

Mouse models genetically engineered to express human genes associated with neurodegenerative diseases, including AD, have significantly advanced our understanding of the underlying pathogenic mechanisms driving these conditions. However, the relative resistance to neurotoxicity observed in these mice has puzzled scientists for decades [87]. As laboratory mice have relatively long telomeres and short lifespans, it could be hypothesized that they are particularly resistant to certain forms of senescence. Specifically, laboratory mice only live 18 to 24 months and present telomeres that are 25–150 kb long at birth, while human telomeres are around 10-15 kb long [88, 89]. Our results demonstrate that a pathological aging context, modeled here by inducing senescence through telomere damage, may be required to trigger extensive brain atrophy and neuronal loss in transgenic mice overexpressing aggregation-prone human tau. Consequently, this combined model of senescence and tauopathy could serve as a valuable preclinical research model to more accurately recapitulate the extent of neuronal death observed in tau-related human diseases and evaluate potential therapeutic targets.

## Conclusion

Aging is strongly associated with neurodegenerative diseases, yet an ongoing debate persists regarding whether senescence is a cause or consequence of their onset and progression. Results from our study proved that senescence acts as an upstream modulator of tauopathy, driving tau hyperphosphorylation and aggregation, which in turn promotes neuroinflammation and neurodegeneration. Taken together, our data highlights the critical need to study brain senescence as a key step towards identifying therapeutic targets for AD and other neurodegenerative diseases cursing with tauopathy.

## List of abbreviations

AD: Alzheimer’s disease
APP: amyloid precursor protein
Aβ: amyloid β
CA: cornu Ammonis
DG: dentate gyrus
ND: neurodegenerative disease
NFT: neurofibrillary tangles
SASP: senescence-associated secretory phenotype
WB: Western blot
WT: wild-type

## Declarations

### Ethics approval and consent to participate

All animal procedures were carried out in compliance with institutional and European guidelines and were approved by the Ethical Committee for Animal Welfare at UCLouvain (References: 2018/UCL/MD/38 and 2021/UCL/MD/18).

### Consent for publication

Not applicable

### Availability of data and materials

All datasets generated and analyzed during this study are included in this published article and its supplementary material.

### Authors’ contributions

DP and NS designed and performed research, analyzed and interpreted data and wrote the manuscript. AV and JG performed experiments and analyzed and interpreted data. NS and PKC conceived, designed and supervised the research project. All authors reviewed and approved the final version of the manuscript.

## Acknowledgements

We thank the team of Peter Davies for kindly providing the PHF-1 antibody.

## References

1. Hou Y, Dan X, Babbar M, Wei Y, Hasselbalch SG, Croteau DL, Bohr VA: Ageing as a risk factor for neurodegenerative disease. Nat Rev Neurol 2019, 15:565–581.

2. Gustavsson A, Norton N, Fast T, Frölich L, Georges J, Holzapfel D, Kirabali T, Krolak-Salmon P, Rossini PM, Ferretti MT, et al: Global estimates on the number of persons across the Alzheimer’s disease continuum. Alzheimers Dement 2023, 19:658–670.

3. GBD 2019 Dementia Forecasting Collaborators: Estimation of the global prevalence of dementia in 2019 and forecasted prevalence in 2050: an analysis for the Global Burden of Disease Study 2019. Lancet Public Health 2022, 7:e105–e125.

4. Campisi J, d’Adda di Fagagna F: Cellular senescence: when bad things happen to good cells. Nat Rev Mol Cell Biol 2007, 8:729–740.

5. Lushchak O, Schosserer M, Grillari J: Senopathies-Diseases Associated with Cellular Senescence. Biomolecules 2023, 13.

6. Götz J, Halliday G, Nisbet RM: Molecular Pathogenesis of the Tauopathies. Annu Rev Pathol 2019, 14:239–261.

7. Hardy J: Alzheimer’s disease: the amyloid cascade hypothesis: an update and reappraisal. J Alzheimers Dis 2006, 9:151–153.

8. Arriagada PV, Growdon JH, Hedley-Whyte ET, Hyman BT: Neurofibrillary tangles but not senile plaques parallel duration and severity of Alzheimer’s disease. Neurology 1992, 42:631–639.

9. Nelson PT, Alafuzoff I, Bigio EH, Bouras C, Braak H, Cairns NJ, Castellani RJ, Crain BJ, Davies P, Del Tredici K, et al: Correlation of Alzheimer disease neuropathologic changes with cognitive status: a review of the literature. J Neuropathol Exp Neurol 2012, 71:362–381.

10. Zhang Y, Chen H, Li R, Sterling K, Song W: Amyloid β-based therapy for Alzheimer’s disease: challenges, successes and future. Signal Transduct Target Ther 2023, 8:248.

11. Musi N, Valentine JM, Sickora KR, Baeuerle E, Thompson CS, Shen Q, Orr ME: Tau protein aggregation is associated with cellular senescence in the brain. Aging Cell 2018, 17:e12840.

12. Herdy JR, Traxler L, Agarwal RK, Karbacher L, Schlachetzki JCM, Boehnke L, Zangwill D, Galasko D, Glass CK, Mertens J, Gage FH: Increased post-mitotic senescence in aged human neurons is a pathological feature of Alzheimer’s disease. Cell Stem Cell 2022, 29:1637–1652.e1636.

13. Hu Y, Fryatt GL, Ghorbani M, Obst J, Menassa DA, Martin-Estebane M, Muntslag TAO, Olmos-Alonso A, Guerrero-Carrasco M, Thomas D, et al: Replicative senescence dictates the emergence of disease-associated microglia and contributes to Aβ pathology. Cell Rep 2021, 35:109228.

14. Bhat R, Crowe EP, Bitto A, Moh M, Katsetos CD, Garcia FU, Johnson FB, Trojanowski JQ, Sell C, Torres C: Astrocyte senescence as a component of Alzheimer’s disease. PLoS One 2012, 7:e45069.

15. Gaikwad S, Puangmalai N, Bittar A, Montalbano M, Garcia S, McAllen S, Bhatt N, Sonawane M, Sengupta U, Kayed R: Tau oligomer induced HMGB1 release contributes to cellular senescence and neuropathology linked to Alzheimer’s disease and frontotemporal dementia. Cell Rep 2021, 36:109419.

16. Dehkordi SK, Walker J, Sah E, Bennett E, Atrian F, Frost B, Woost B, Bennett RE, Orr TC, Zhou Y, et al: Profiling senescent cells in human brains reveals neurons with CDKN2D/p19 and tau neuropathology. Nat Aging 2021, 1:1107–1116.

17. Fancy NN, Smith AM, Caramello A, Tsartsalis S, Davey K, Muirhead RCJ, McGarry A, Jenkyns MH, Schneegans E, Chau V, et al: Characterisation of premature cell senescence in Alzheimer’s disease using single nuclear transcriptomics. Acta Neuropathol 2024, 147:78.

18. Bussian TJ, Aziz A, Meyer CF, Swenson BL, van Deursen JM, Baker DJ: Clearance of senescent glial cells prevents tau-dependent pathology and cognitive decline. Nature 2018, 562:578–582.

19. Wei Z, Chen XC, Song Y, Pan XD, Dai XM, Zhang J, Cui XL, Wu XL, Zhu YG: Amyloid β Protein Aggravates Neuronal Senescence and Cognitive Deficits in 5XFAD Mouse Model of Alzheimer’s Disease. Chin Med J (Engl) 2016, 129:1835–1844.

20. Forero DA, González-Giraldo Y, López-Quintero C, Castro-Vega LJ, Barreto GE, Perry G: Meta-analysis of Telomere Length in Alzheimer’s Disease. J Gerontol A Biol Sci Med Sci 2016, 71:1069–1073.

21. Gao K, Wei C, Zhu J, Wang X, Chen G, Luo Y, Zhang D, Yue W, Yu H: Frontiers | Exploring the Causal Pathway From Telomere Length to Alzheimer’s Disease: An Update Mendelian Randomization Study. Frontiers in Psychiatry 2019/11/15, 10.

22. Scarabino D, Broggio E, Gambina G, Pelliccia F, Corbo RM: Common variants of human TERT and TERC genes and susceptibility to sporadic Alzheimers disease. Exp Gerontol 2017, 88:19–24.

23. Zhan Y, Song C, Karlsson R, Tillander A, Reynolds CA, Pedersen NL, Hägg S: Telomere Length Shortening and Alzheimer Disease--A Mendelian Randomization Study. JAMA Neurol 2015, 72:1202–1203.

24. Saez-Atienzar S, Masliah E: Cellular senescence and Alzheimer disease: the egg and the chicken scenario. Nat Rev Neurosci 2020, 21:433–444.

25. Blasco MA, Lee HW, Hande MP, Samper E, Lansdorp PM, DePinho RA, Greider CW: Telomere shortening and tumor formation by mouse cells lacking telomerase RNA. Cell 1997, 91:25–34.

26. Yoshiyama Y, Higuchi M, Zhang B, Huang SM, Iwata N, Saido TC, Maeda J, Suhara T, Trojanowski JQ, Lee VM: Synapse loss and microglial activation precede tangles in a P301S tauopathy mouse model. Neuron 2007, 53:337–351.

27. Suelves N, Saleki S, Ibrahim T, Palomares D, Moonen S, Koper MJ, Vrancx C, Vadukul DM, Papadopoulos N, Viceconte N, et al: Senescence-related impairment of autophagy induces toxic intraneuronal amyloid-β accumulation in a mouse model of amyloid pathology. Acta Neuropathol Commun 2023, 11:82.

28. Cawthon RM: Telomere measurement by quantitative PCR. Nucleic Acids Res 2002, 30:e47.

29. Schneider CA, Rasband WS, Eliceiri KW: NIH Image to ImageJ: 25 years of image analysis. Nature Methods 2012, 9:671–675.

30. Gf P, Franklin K: The Mouse Brain In Stereotaxic Coordinates. 2003.

31. Baumann P, Schriever SC, Kullmann S, Zimprich A, Feuchtinger A, Amarie O, Peter A, Walch A, Gailus-Durner V, Fuchs H, et al: Dusp8 affects hippocampal size and behavior in mice and humans. Scientific Reports 2019, 9:19483.

32. Rogakou EP, Pilch DR, Orr AH, Ivanova VS, Bonner WM: DNA double-stranded breaks induce histone H2AX phosphorylation on serine 139. J Biol Chem 1998, 273:5858–5868.

33. Maruyama M, Shimada H, Suhara T, Shinotoh H, Ji B, Maeda J, Zhang MR, Trojanowski JQ, Lee VM, Ono M, et al: Imaging of tau pathology in a tauopathy mouse model and in Alzheimer patients compared to normal controls. Neuron 2013, 79:1094–1108.

34. Goedert M, Jakes R, Vanmechelen E: Monoclonal antibody AT8 recognises tau protein phosphorylated at both serine 202 and threonine 205. Neurosci Lett 1995, 189:167–169.

35. Braak H, Alafuzoff I, Arzberger T, Kretzschmar H, Del Tredici K: Staging of Alzheimer disease-associated neurofibrillary pathology using paraffin sections and immunocytochemistry. Acta Neuropathol 2006, 112:389–404.

36. Basheer N, Smolek T, Hassan I, Liu F, Iqbal K, Zilka N, Novak P: Does modulation of tau hyperphosphorylation represent a reasonable therapeutic strategy for Alzheimer’s disease? From preclinical studies to the clinical trials. Molecular Psychiatry 2023, 28:2197–2214.

37. Liu C, Götz J: Profiling murine tau with 0N, 1N and 2N isoform-specific antibodies in brain and peripheral organs reveals distinct subcellular localization, with the 1N isoform being enriched in the nucleus. PLoS One 2013, 8:e84849.

38. McMillan P, Korvatska E, Poorkaj P, Evstafjeva Z, Robinson L, Greenup L, Leverenz J, Schellenberg GD, D’Souza I: Tau isoform regulation is region-and cell-specific in mouse brain. Journal of Comparative Neurology 2008, 511:788–803.

39. Petry FR, Pelletier J, Bretteville A, Morin F, Calon F, Hébert SS, Whittington RA, Planel E: Specificity of anti-tau antibodies when analyzing mice models of Alzheimer’s disease: problems and solutions. PLoS One 2014, 9:e94251.

40. Yang T, Liu H, Tran KC, Leng A, Massa SM, Longo FM: Small-molecule modulation of the p75 neurotrophin receptor inhibits a wide range of tau molecular pathologies and their sequelae in P301S tauopathy mice. Acta Neuropathol Commun 2020, 8:156.

41. Ellis MJ, Lekka C, Holden KL, Tulmin H, Seedat F, O’Brien DP, Dhayal S, Zeissler ML, Knudsen JG, Kessler BM, et al: Identification of high-performing antibodies for the reliable detection of Tau proteoforms by Western blotting and immunohistochemistry. Acta Neuropathol 2024, 147:87.

42. Zilka N, Filipcik P, Koson P, Fialova L, Skrabana R, Zilkova M, Rolkova G, Kontsekova E, Novak M: Truncated tau from sporadic Alzheimer’s disease suffices to drive neurofibrillary degeneration in vivo. FEBS Lett 2006, 580:3582–3588.

43. Gu J, Xu W, Jin N, Li L, Zhou Y, Chu D, Gong CX, Iqbal K, Liu F: Truncation of Tau selectively facilitates its pathological activities. J Biol Chem 2020, 295:13812–13828.

44. Zhang W, Xiao D, Mao Q, Xia H: Role of neuroinflammation in neurodegeneration development. Signal Transduction and Targeted Therapy 2023, 8:267.

45. Heneka MT, van der Flier WM, Jessen F, Hoozemanns J, Thal DR, Boche D, Brosseron F, Teunissen C, Zetterberg H, Jacobs AH, et al: Neuroinflammation in Alzheimer disease. Nature Reviews Immunology 2025, 25:321–352.

46. Von Bernhardi R, Eugenin-von Bernhardi L, Eugenin J: Microglial cell dysregulation in Brain Aging and Neurodegeneration. Frontiers in Aging Neuroscience 2015, Volume 7 - 2015.

47. Streit WJ, Braak H, Xue QS, Bechmann I: Dystrophic (senescent) rather than activated microglial cells are associated with tau pathology and likely precede neurodegeneration in Alzheimer’s disease. Acta Neuropathol 2009, 118:475–485.

48. Gildea HK, Liddelow SA: Mechanisms of astrocyte aging in reactivity and disease. Mol Neurodegener 2025, 20:21.

49. Bachstetter AD, Van Eldik LJ, Schmitt FA, Neltner JH, Ighodaro ET, Webster SJ, Patel E, Abner EL, Kryscio RJ, Nelson PT: Disease-related microglia heterogeneity in the hippocampus of Alzheimer’s disease, dementia with Lewy bodies, and hippocampal sclerosis of aging. Acta Neuropathologica Communications 2015, 3:32.

50. Tzioras M, McGeachan RI, Durrant CS, Spires-Jones TL: Synaptic degeneration in Alzheimer disease. Nature Reviews Neurology 2023, 19:19–38.

51. Wang C, Xiong M, Gratuze M, Bao X, Shi Y, Andhey PS, Manis M, Schroeder C, Yin Z, Madore C, et al: Selective removal of astrocytic APOE4 strongly protects against tau-mediated neurodegeneration and decreases synaptic phagocytosis by microglia. Neuron 2021, 109:1657–1674.e1657.

52. Vogels T, Murgoci A-N, Hromádka T: Intersection of pathological tau and microglia at the synapse. Acta Neuropathologica Communications 2019, 7:109.

53. Zhu J, Wu C, Yang L: Cellular senescence in Alzheimer’s disease: from physiology to pathology. Transl Neurodegener 2024, 13:55.

54. Shanbhag NM, Evans MD, Mao W, Nana AL, Seeley WW, Adame A, Rissman RA, Masliah E, Mucke L: Early neuronal accumulation of DNA double strand breaks in Alzheimer’s disease. Acta Neuropathologica Communications 2019, 7:77.

55. Li LJ, Sun XY, Huang YR, Lu S, Xu YM, Yang J, Xie XX, Zhu J, Niu XY, Wang D, et al: Neuronal double-stranded DNA accumulation induced by DNase II deficiency drives tau phosphorylation and neurodegeneration. Transl Neurodegener 2024, 13:39.

56. Li LJ, Liang SY, Sun XY, Zhu J, Niu XY, Du XY, Huang YR, Liu RT: Microglial double stranded DNA accumulation induced by DNase II deficiency drives neuroinflammation and neurodegeneration. J Neuroinflammation 2025, 22:11.

57. Lasagna-Reeves CA, Castillo-Carranza DL, Sengupta U, Clos AL, Jackson GR, Kayed R: Tau oligomers impair memory and induce synaptic and mitochondrial dysfunction in wild-type mice. Molecular Neurodegeneration 2011, 6:39.

58. Patterson KR, Remmers C, Fu Y, Brooker S, Kanaan NM, Vana L, Ward S, Reyes JF, Philibert K, Glucksman MJ, Binder LI: Characterization of Prefibrillar Tau Oligomers in Vitro and in Alzheimer Disease. Journal of Biological Chemistry 2011, 286:23063–23076.

59. Pei JJ, Gong CX, An WL, Winblad B, Cowburn RF, Grundke-Iqbal I, Iqbal K: Okadaic-acid-induced inhibition of protein phosphatase 2A produces activation of mitogen-activated protein kinases ERK1/2, MEK1/2, and p70 S6, similar to that in Alzheimer’s disease. Am J Pathol 2003, 163:845-858.

60. Ksiezak-Reding H, Pyo HK, Feinstein B, Pasinetti GM: Akt/PKB kinase phosphorylates separately Thr212 and Ser214 of tau protein in vitro. Biochim Biophys Acta 2003, 1639:159–168.

61. Silva MC, Nandi GA, Tentarelli S, Gurrell IK, Jamier T, Lucente D, Dickerson BC, Brown DG, Brandon NJ, Haggarty SJ: Prolonged tau clearance and stress vulnerability rescue by pharmacological activation of autophagy in tauopathy neurons. Nature Communications 2020, 11:3258.

62. Chesser AS, Pritchard SM, Johnson GV: Tau clearance mechanisms and their possible role in the pathogenesis of Alzheimer disease. Front Neurol 2013, 4:122.

63. Lee MJ, Lee JH, Rubinsztein DC: Tau degradation: The ubiquitin–proteasome system versus the autophagy-lysosome system. Progress in Neurobiology 2013, 105:49–59.

64. Hamano T, Enomoto S, Shirafuji N, Ikawa M, Yamamura O, Yen S-H, Nakamoto Y: Autophagy and Tau Protein. International Journal of Molecular Sciences 2021, 22:7475.

65. Kang Y, Zhang H, Zhao Y, Wang Y, Wang W, He Y, Zhang W, Zhang W, Zhu X, Zhou Y, et al: Telomere Dysfunction Disturbs Macrophage Mitochondrial Metabolism and the NLRP3 Inflammasome through the PGC-1α/TNFAIP3 Axis. Cell Rep 2018, 22:3493–3506.

66. Bartolome F, Carro E, Alquezar C: Oxidative Stress in Tauopathies: From Cause to Therapy. Antioxidants (Basel) 2022, 11.

67. Samluk L, Ostapczuk P, Dziembowska M: Long-term mitochondrial stress induces early steps of Tau aggregation by increasing reactive oxygen species levels and affecting cellular proteostasis. Mol Biol Cell 2022, 33:ar67.

68. Su B, Wang X, Lee HG, Tabaton M, Perry G, Smith MA, Zhu X: Chronic oxidative stress causes increased tau phosphorylation in M17 neuroblastoma cells. Neurosci Lett 2010, 468:267–271.

69. Melov S, Adlard PA, Morten K, Johnson F, Golden TR, Hinerfeld D, Schilling B, Mavros C, Masters CL, Volitakis I, et al: Mitochondrial oxidative stress causes hyperphosphorylation of tau. PLoS One 2007, 2:e536.

70. Rissman RA, Poon WW, Blurton-Jones M, Oddo S, Torp R, Vitek MP, LaFerla FM, Rohn TT, Cotman CW: Caspase-cleavage of tau is an early event in Alzheimer disease tangle pathology. J Clin Invest 2004, 114:121–130.

71. Rohn TT, Rissman RA, Davis MC, Kim YE, Cotman CW, Head E: Caspase-9 activation and caspase cleavage of tau in the Alzheimer’s disease brain. Neurobiol Dis 2002, 11:341–354.

72. Wray S, Saxton M, Anderton BH, Hanger DP: Direct analysis of tau from PSP brain identifies new phosphorylation sites and a major fragment of N-terminally cleaved tau containing four microtubule-binding repeats. J Neurochem 2008, 105:2343–2352.

73. Chen HH, Liu P, Auger P, Lee SH, Adolfsson O, Rey-Bellet L, Lafrance-Vanasse J, Friedman BA, Pihlgren M, Muhs A, et al: Calpain-mediated tau fragmentation is altered in Alzheimer’s disease progression. Sci Rep 2018, 8:16725.

74. Le L, Lee J, Im D, Park S, Hwang KD, Lee JH, Jiang Y, Lee YS, Suh YH, Kim HI, Lee MJ: Self-Aggregating Tau Fragments Recapitulate Pathologic Phenotypes and Neurotoxicity of Alzheimer’s Disease in Mice. Adv Sci (Weinh) 2023, 10:e2302035.

75. Kim Y, Choi H, Lee W, Park H, Kam TI, Hong SH, Nah J, Jung S, Shin B, Lee H, et al: Caspase-cleaved tau exhibits rapid memory impairment associated with tau oligomers in a transgenic mouse model. Neurobiol Dis 2016, 87:19–28.

76. Gao Y, Wang Y, Lei H, Xu Z, Li S, Yu H, Xie J, Zhang Z, Liu G, Zhang Y, et al: A novel transgenic mouse line with hippocampus-dominant and inducible expression of truncated human tau. Transl Neurodegener 2023, 12:51.

77. Wang YP, Biernat J, Pickhardt M, Mandelkow E, Mandelkow EM: Stepwise proteolysis liberates tau fragments that nucleate the Alzheimer-like aggregation of full-length tau in a neuronal cell model. Proc Natl Acad Sci U S A 2007, 104:10252–10257.

78. Chung CW, Song YH, Kim IK, Yoon WJ, Ryu BR, Jo DG, Woo HN, Kwon YK, Kim HH, Gwag BJ, et al: Proapoptotic effects of tau cleavage product generated by caspase-3. Neurobiol Dis 2001, 8:162–172.

79. Sedarous M, Keramaris E, O’Hare M, Melloni E, Slack RS, Elce JS, Greer PA, Park DS: Calpains mediate p53 activation and neuronal death evoked by DNA damage. J Biol Chem 2003, 278:26031–26038.

80. Vakifahmetoglu H, Olsson M, Orrenius S, Zhivotovsky B: Functional connection between p53 and caspase-2 is essential for apoptosis induced by DNA damage. Oncogene 2006, 25:5683–5692.

81. Braak H, Thal DR, Ghebremedhin E, Del Tredici K: Stages of the Pathologic Process in Alzheimer Disease: Age Categories From 1 to 100 Years. Journal of Neuropathology & Experimental Neurology 2011, 70:960–969.

82. Mrdjen D, Fox EJ, Bukhari SA, Montine KS, Bendall SC, Montine TJ: The basis of cellular and regional vulnerability in Alzheimer’s disease. Acta Neuropathologica 2019, 138:729–749.

83. Laurent C, Buée L, Blum D: Tau and neuroinflammation: What impact for Alzheimer’s Disease and Tauopathies? Biomed J 2018, 41:21–33.

84. Khan AM, Babcock AA, Saeed H, Myhre CL, Kassem M, Finsen B: Telomere dysfunction reduces microglial numbers without fully inducing an aging phenotype. Neurobiol Aging 2015, 36:2164–2175.

85. Raj DD, Moser J, van der Pol SM, van Os RP, Holtman IR, Brouwer N, Oeseburg H, Schaafsma W, Wesseling EM, den Dunnen W, et al: Enhanced microglial pro-inflammatory response to lipopolysaccharide correlates with brain infiltration and blood-brain barrier dysregulation in a mouse model of telomere shortening. Aging Cell 2015, 14:1003–1013.

86. Scheffold A, Holtman IR, Dieni S, Brouwer N, Katz SF, Jebaraj BM, Kahle PJ, Hengerer B, Lechel A, Stilgenbauer S, et al: Telomere shortening leads to an acceleration of synucleinopathy and impaired microglia response in a genetic mouse model. Acta Neuropathol Commun 2016, 4:87.

87. Dawson TM, Golde TE, Lagier-Tourenne C: Animal models of neurodegenerative diseases. Nat Neurosci 2018, 21:1370–1379.

88. Kakuo S, Asaoka K, Ide T: Human is a unique species among primates in terms of telomere length. Biochem Biophys Res Commun 1999, 263:308–314.

89. Gomes NM, Ryder OA, Houck ML, Charter SJ, Walker W, Forsyth NR, Austad SN, Venditti C, Pagel M, Shay JW, Wright WE: Comparative biology of mammalian telomeres: hypotheses on ancestral states and the roles of telomeres in longevity determination. Aging Cell 2011, 10:761–768.

