## Supplementary Information for "Telomere-driven senescence accelerates tau pathology, neuroinflammation and neurodegeneration in a tauopathy mouse model"

### Supplementary Figures and Legends

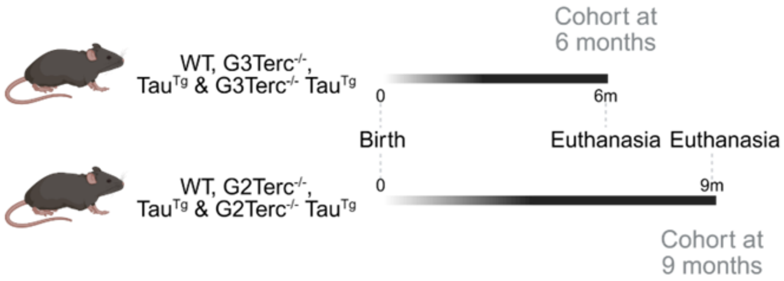


**Figure S1. Schematic representation of the two senescence models and experimental time points used for our study.** WT, G3Terc$^{-/-}$, Tau$^{Tg}$ and G3Terc$^{-/-}$ Tau$^{Tg}$ mice were sacrificed, and their brains collected at 6 months. WT, G2Terc$^{-/-}$, Tau$^{Tg}$ and G2Terc$^{-/-}$ Tau$^{Tg}$ mice were sacrificed, and their brains collected at 9 months. Created in BioRender. Kienlen-Campard, P. (2025) https://BioRender.com/k4g060o.

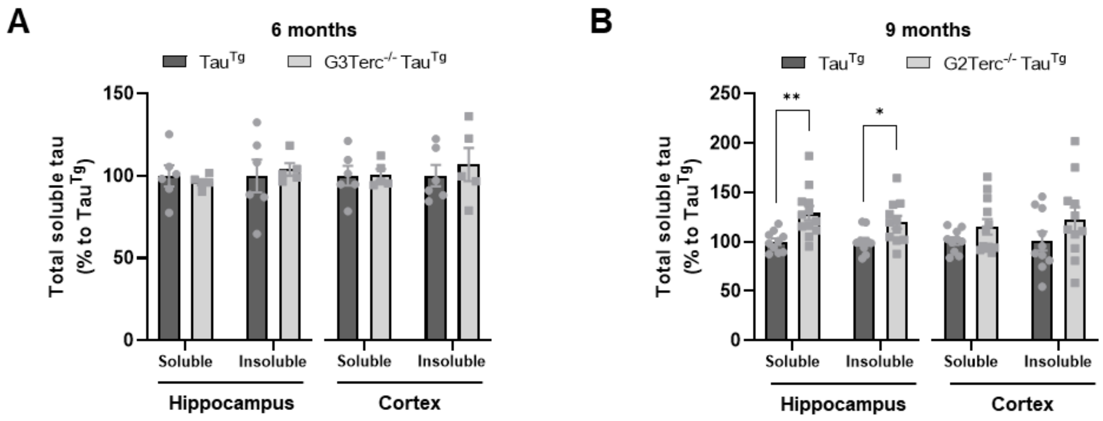


**Figure S2. Total tau levels are significantly increased in a senescent-tauopathy context in relatively aged mice.** Quantification of total tau levels after normalization to actin (Figure 3) in hippocampal and cortical protein extracts of 6-month-old Tau$^{Tg}$ and G3Terc$^{-/-}$ Tau$^{Tg}$ (**A**) and 9-month-old Tau$^{Tg}$ and G2Terc$^{-/-}$ Tau$^{Tg}$ mice (**B**). *$P < 0.05$, **$P < 0.01$ (Student's *t*-test or Mann-Whitney's test, n= 5-12 mice/group).

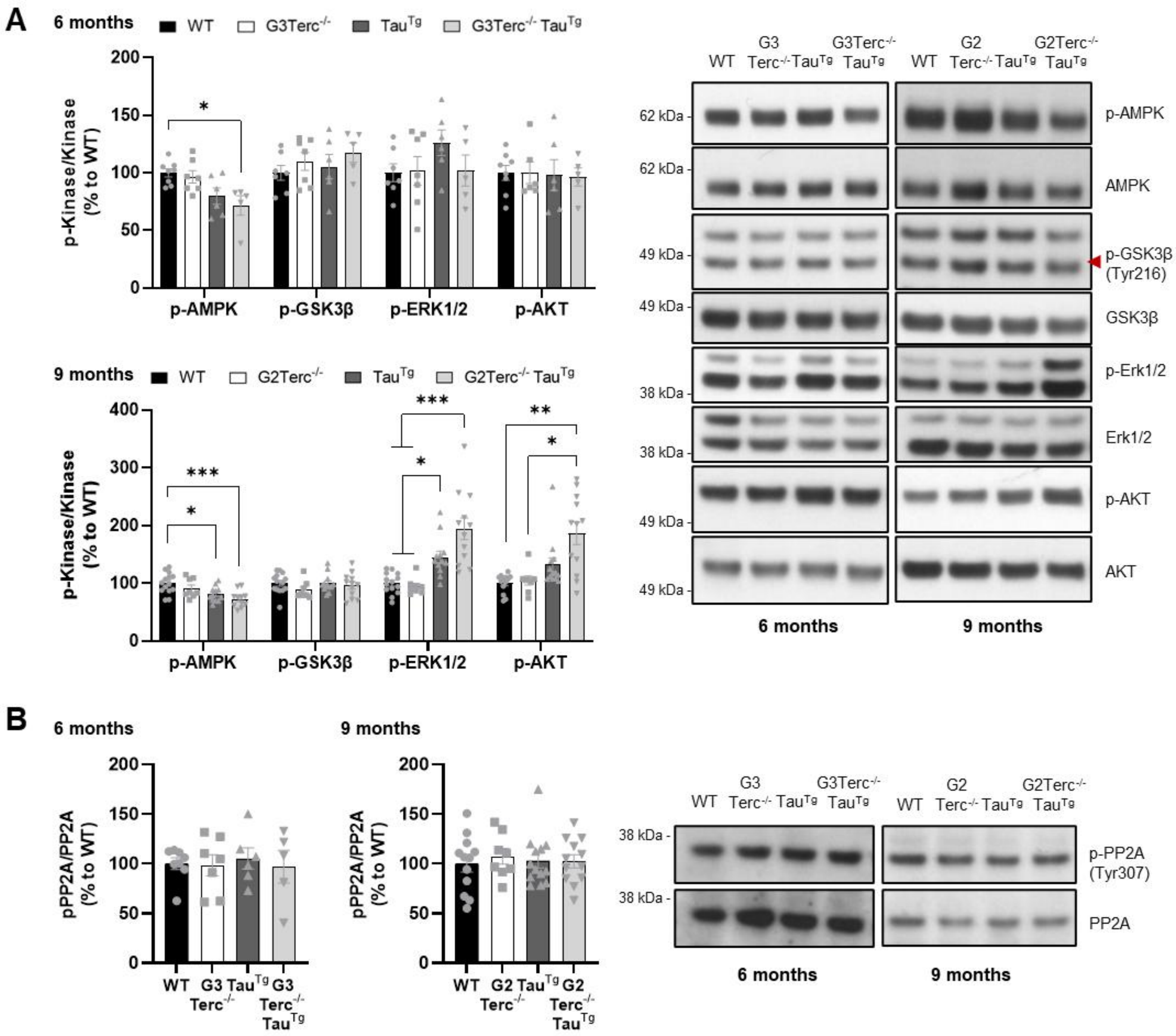


**Figure S3. Telomere-induced senescence does not modify protein levels of tau kinases and phosphatases, nor their phosphorylated status at early stages of the pathology.** WB evaluation of major tau kinases (**A**) and the main tau phosphatase PP2A (**B**) in hippocampal protein extracts from 6-month-old WT, G3Terc$^{-/-}$, Tau$^{Tg}$ and G3Terc$^{-/-}$ Tau$^{Tg}$, and 9-month-old WT, G2Terc$^{-/-}$, Tau$^{Tg}$ and G2Terc$^{-/-}$ Tau$^{Tg}$ mice. *$P < 0.05$, **$P < 0.01$, **$P < 0.001$ (One-way ANOVA with Tukey's post-hoc analysis or Kruskal-Wallis test, n= 5-13 mice/group).

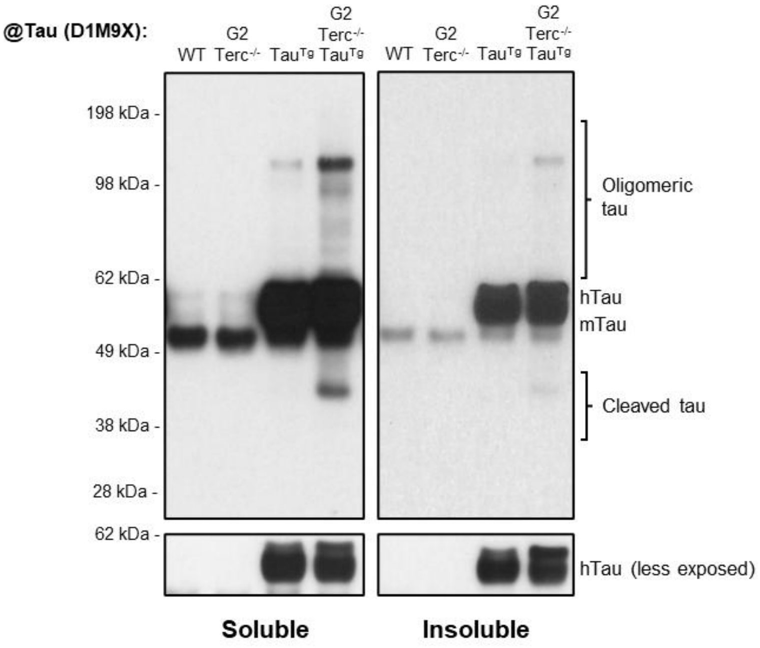


**Figure S4. Lack of truncated and oligomeric tau in non-tauopathy mice.** WB analysis of oligomeric and truncated Tau levels in soluble and insoluble protein extracts from 9-month-old WT, G2Terc$^{-/-}$, Tau$^{Tg}$ and G2Terc$^{-/-}$ Tau$^{Tg}$ mouse hippocampi, using a total tau antibody (D1M9X clone). In the absence of mutated human tau (tauopathy conditions), there are no visible bands corresponding to truncated and oligomeric tau.

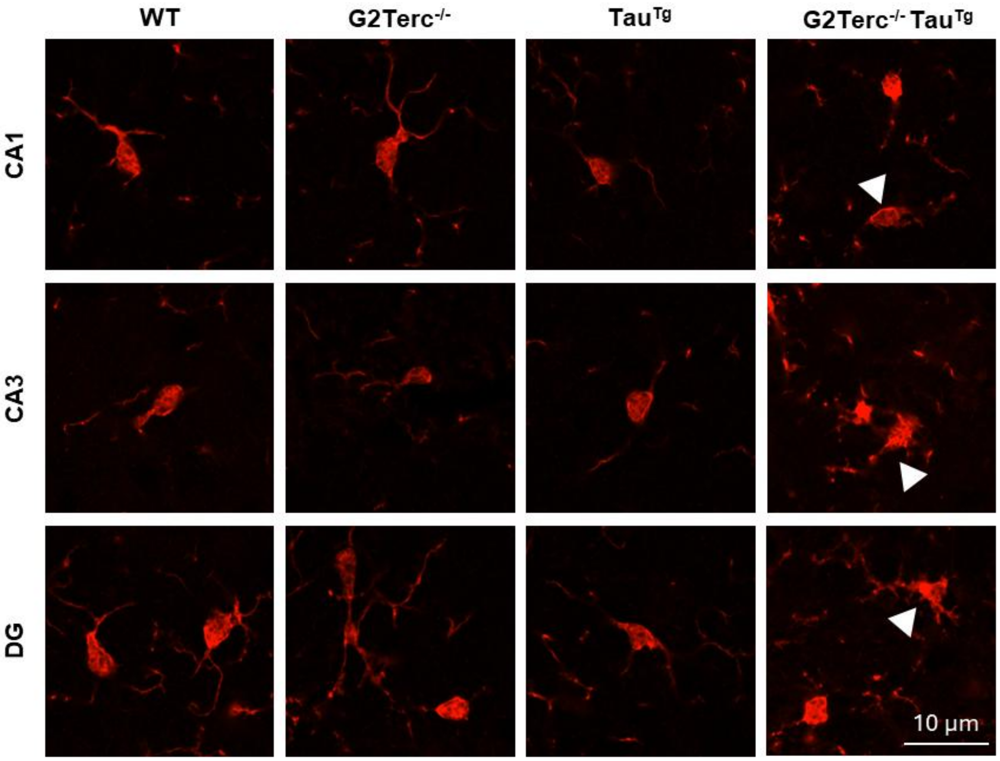


**Figure S5. Dystrophic microglia is observed in G2Terc$^{-/-}$ Tau$^{tg}$ brains.** Iba1 (red) immunofluorescence representative images of 9-month-old WT, G2Terc$^{-/-}$, Tau$^{Tg}$ and G2Terc$^{-/-}$ Tau$^{Tg}$ brains by confocal microscopy. Apparent beading and fragmentation of microglia projections, signs of dystrophia, were observed exclusively in the G2Terc$^{-/-}$ Tau$^{Tg}$ condition.

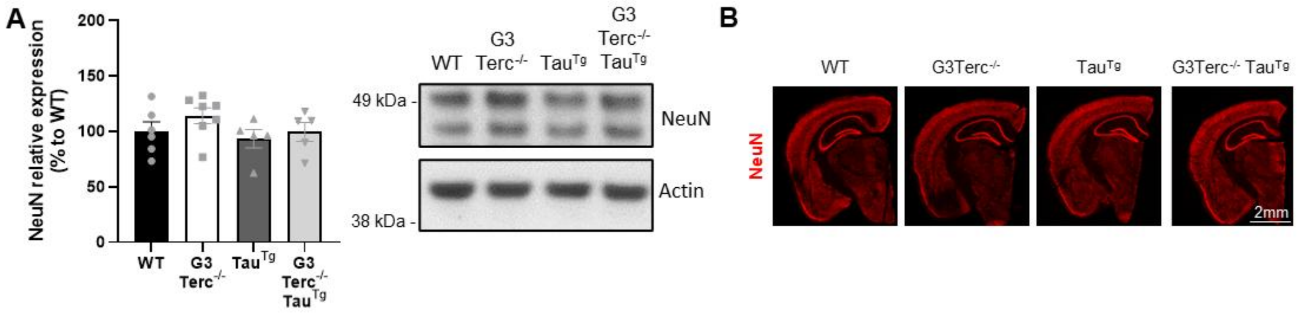


**Figure S6. Senescence does not affect neurodegeneration in G3Terc$^{-/-}$ Tau$^{Tg}$ brains at 6 months. A)** WB evaluation of NeuN expression in hippocampal extracts from 6-month-old WT, G3Terc$^{-/-}$, Tau$^{Tg}$ and G3Terc$^{-/-}$ Tau$^{Tg}$ mice. One-way ANOVA with Tukey's post-hoc analysis, n= 5-6 mice/group. **B**) NeuN immunofluorescence representative images from WT, G3Terc$^{-/-}$, Tau$^{Tg}$ and G3Terc$^{-/-}$ Tau$^{Tg}$ mice.

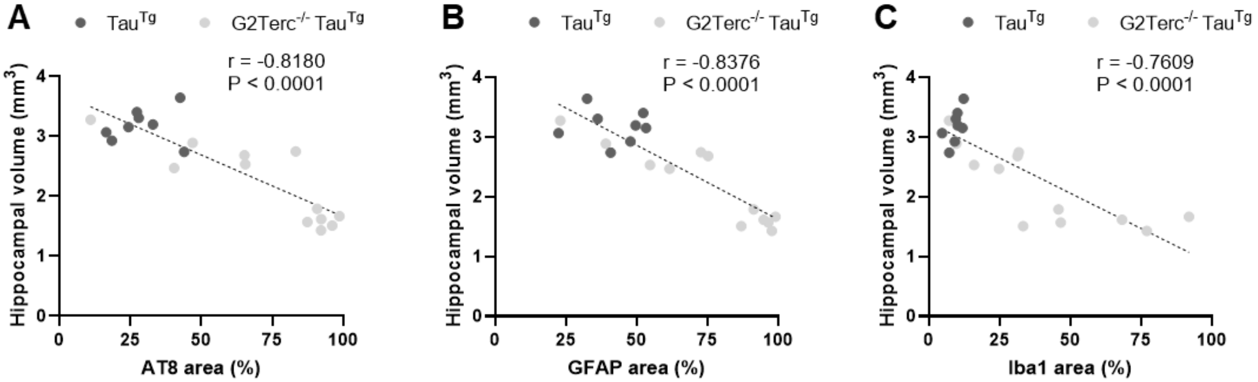


**Figure S7. Pathological tau immunostaining and neuroinflammation negatively correlate with hippocampal volume in G2Terc$^{-/-}$ Tau$^{Tg}$ mice**. **A**) Correlation analysis of the relationship between hyperphosphorylated tau (AT8-positive area) and hippocampal volume (mm$^3$) in 9-month-old Tau$^{Tg}$ and G2Terc$^{-/-}$ Tau$^{Tg}$ mice (Figures 2 and 7, Spearman correlation, n=20). **B-C**) Correlation analysis of the relationship between hyperphosphorylated tau (AT8 area) and neuroinflammatory markers of astrogliosis (GFAP area) (**B**) and microgliosis (Iba1 area) (**C**) in 9-month-old Tau$^{Tg}$ and G2Terc$^{-/-}$ Tau$^{Tg}$ mice (Figures 2 and 5, Spearman correlation, n=20).

| Target | Forward (5'- 3') | Reverse (5'- 3') |
|---|---|---|
| Il1b | GCAACTGTTCCTGAACTCAACT | ATCTTTTGGGGTCCGTCAACT |
| Cxcl1 | AACCGAAGTCATAGCCACAC | GACACCTTTTAGCATCTTTTGG |
| p19 | CGCAGGTTCTTGGTCACTGT | TGTTCACAGAAGCCAGAGCG |
| p21 | CCTGGTGATGTCCGACCTG | CCATGAGCGCATCGCAATC |
| Tbp | GATGTGCGTCAGGCGTTC | GGGTTATCTTCACACACCATGA |
| Gapdh | ACCCAGAAGACTGTGGATGG | ACACATTGGGGGTAGGAAC |

**Table S1. Sequences of primers used for quantitative RT-PCR.**

| Target | Clone | Source | Reference | Application (Dilution) |
|---|---|---|---|---|
| Actin | Polyclonal | Sigma-Aldrich | A2066 | WB (1:5,000) |
| AMPKα | Polyclonal | Cell Signaling | 2532 | WB (1:1,000) |
| CDK5 | J-3 | Santa Cruz Biotechnology | sc-6247 | WB (1:500) |
| Erk1/2 | 137F5 | Cell Signaling | 4695 | WB (1:1,000) |
| GFAP | ASTRO6 | Thermo Fisher Scientific | MA5-12023 | WB (1:1,000) |
| GFAP | Polyclonal | Abcam | ab4674 | IF (1:1,000) |
| GluR1 | C3T | Sigma-Aldrich | 04-855 | WB (1:5,000) |
| GSK3β | 27C10 | Cell Signaling | 9315 | WB (1:2,000) |
| Iba1 | EPR16588 | Abcam | ab178846 | IF (1:500), WB (1:500) |
| NeuN | Polyclonal | Abcam | ab104225 | IF (1:1,000), WB (1:10,000) |
| NMDAR2B | 13/NMDAR2B | BD Biosciences | 610417 | WB (1:500) |
| pAMPKα Thr172 | 40H9 | Cell Signaling | 2535 | WB (1:1,000) |
| pErk1/2 Thr202/Tyr204 | (D13.14.4E) XP® | Cell Signaling | 4370 | WB (1:1,000) |
| pGSK3β Tyr216 + pGSK3α Tyr279 | Polyclonal | Abcam | ab75745 | WB (1:1,000) |
| PSD-95 | Polyclonal | Abcam | ab18258 | WB (1:2,000) |
| pTau Ser202 + Thr205 | AT8 | ThermoFisher | MN1020 | IF (1:500), WB (1:3,000) |
| pTau Ser396 + Ser404 | PHF-1 | Peter Davies | / | WB (1:1,000) |
| pTau Ser422 | Polyclonal | ThermoFisher | 44-764G | WB (1:1,000) |
| pTau Thr181 | AT270 | ThermoFisher | MN1050 | WB (1:3,000) |
| pTau Thr231 | AT180 | ThermoFisher | MN1040 | WB (1:1,000) |
| pTau Thr262 | Polyclonal | ThermoFisher | 44-750G | WB (1:1,000) |
| Synaptophysin | Polyclonal | Synaptic Systems | 101 002 | WB (1:50,000) |
| Total tau | D1M9X | Cell Signaling | 46687 | WB (1:3,000) |
| γH2AX | Polyclonal | Cell Signaling | 2577 | WB (1:1,000) |

**Table S2. Information on primary antibodies**
